# A multi-scale forecasting framework to prioritize clinically-relevant antibiotic resistance alleles

**DOI:** 10.64898/2026.11.20.768016

**Authors:** Nancy Obeng, Anna Rodoni, Quirin Niggli, David Dylus, Kenneth Bradley, Alan James Mueller-Breckenridge, Andrej Trauner, Justin Pritchard

## Abstract

Established *in vitro* measures for the emergence of antimicrobial resistance, a critical global threat, can fail to predict which *in vitro* discovered mutations will dominate in the real world following widespread antibiotic use. Here, we introduce a multi-scale computational framework integrating genetic constraints and population pharmacokinetics and apply it to explain resistance allele success in *Acinetobacter baumannii* for the historic case of resistance to the fluoroquinolone antibiotic ciprofloxacin. By coupling intrinsic nucleotide substitution likelihoods with continuous-time stochastic modeling under fluctuating drug exposures, our framework clearly prioritizes the most common real world *gyrA* fluoroquinolone resistance allele S81L over rarer alleles, reflecting its clinical success. While *gyrA* serves as a proof-of-concept case study for direct, active-site drug-target interactions where the target space is constrained, we observe that the nucleotide substitution likelihoods are not explaining allele success in resistance-determining genes *pmrB* and *adeS*. Furthermore, “virtual twin” cohort simulations reveal that restricted drug penetration or compromised immunity expands selective windows, rescuing rare, low-fitness alleles. This advances our understanding of prospective, model-informed risk prediction during antibiotic development.

## Introduction

Antimicrobial resistance is a growing global health threat, already responsible for over a million deaths annually^1^. While it takes decades to develop novel antibiotics, target site and regulatory mutations can lead to high-level resistance within months of clinical deployment^2,3^. This asymmetry between decade-long drug development and rapid mutational adaptation exposes anti-infective pipelines to severe risk of late-stage clinical attrition, loss of potency during treatment and financial devaluation^2–4^. It is thus critical to de-risk the development of novel compounds as early as possible. For effective future and next-generation drugs, discovery and development efforts must therefore anticipate future resistance patterns and move from reactive management of existing resistances to proactive preclinical development targeting potential future resistances^2,5,6^. A key driver of antibiotic resistance are mutations in bacterial genomes, which render treatments ineffective. Thus, we required multi-scale forecasting tools that can map the effect of genome evolution to clinical outcomes in patients^4,7–9^.

To bridge the translational gap between mutations impairing drug efficacy *in vitro* and subsequent treatment success in patients, forecasting tools need to account for the key evolutionary drivers of resistance. Traditional antibiotic drug development heavily relies on retrospective genomic surveillance or phenotypic screens to quantify spontaneous mutation frequencies and identify loci of potential concern^10,11^. While these workflows assess potential for future resistance and help identify genes linked to resistance and nominate potential mutations that might be observed in the clinic, they ignore underlying biochemical, evolutionary and ecological constraints that govern bacterial evolution^12^. While the selective advantage to survive higher drug exposure helps to prioritize which resistance mutation leads to greater minimal inhibitory concentrations (MICs)^10,11^, success of a mutant will depend on the arrival, as well as survival, of the fittest in a given context^5,13–16^. Thus, mutational biases and epistatic constraints that dictate the accessible mutational path to resistance at the molecular level^14,17,18^ are expected to shape which mutants may emerge in patients. Simultaneously, dynamic bottlenecks imposed by the patient environment will determine which variants may survive and expand, and are commonly neglected in current predictive models^19^. During an active infection, bacterial populations do not grow in a vacuum, but are subjected to continuous population turnover driven by drug fluctuations and immune clearance by the host itself^20^. Within-host ecology acts as a profound evolutionary filter layered on the stochastic molecular dynamics of mutation, where even highly beneficial mutations may fail to establish due to random extinction events during early population bottlenecks^21^. Limited drug penetration or pharmacokinetic fluctuations in drug concentration may lead to mutant selection windows that allow less resistant variants to thrive^20,22^. Thus, the ultimate success of a resistance mutant in patients will be determined by evolutionary drivers across scales^5^.

In this paper, we developed a general prospective multi-scale resistance forecasting framework and validated it against *Acinetobacter baumannii*, a critical priority pathogen where rapid resistance evolution severely limits clinical options^23,24^. As a foundation for this predictive platform, we focus on the synthetic fluoroquinolone ciprofloxacin and the resistance target gene *gyrA*, whose evolution is strongly driven by *de novo* point mutations^25^. Retrospective evolutionary analysis of this locus shows that in addition to selection by the drug itself, mutational biases inherent in the *A. baumannii* genome explain the success of the possible resistance alleles and should be leveraged to predict resistance evolution *de novo*. Focusing on direct active-site target modifications in the so-called quinolone resistance-determining region (QRDR), we validate the modeling approach for a case where target space is constrained, as compared to functional pathway modification or efflux derepression via mutation of *pmrB* or *adeS* in colistin or tigecycline resistance, where the impact of mutational bias decays, respectively. Inspired by a computational framework developed for leukemic resistance evolution^5^, we then integrate these multi-scale variables into a continuous-time stochastic model and simulate dynamic population trajectories under fluctuating patient exposures^26^. For *gyrA* mutations, our model accurately forecasts the dominant clinical driver mutation and its clear advantage over rarer alleles. Further, we demonstrate how host-specific factors limiting drug exposure or immune clearance fundamentally shift the balance between mutational supply and competitive establishment across distinct patient cohorts^25^. Ultimately, this forecasting model integrates resistance assessment from descriptive surveillance to prediction, providing a framework to anticipate priority variants and help de-risk antibiotic candidates during preclinical development.

## Results

### Retrospective analysis reveals ultimate drivers of resistance evolution

To identify and quantify key drivers leading to the success of specific resistance alleles, we started with a retrospective analysis of antibiotic resistance evolution. Our study focused on resistance emergence by single nucleotide polymorphisms in resistance-related genes, i.e. drug target genes and related machinery, as these may lead to strong changes in drug susceptibility^17,27,28^. While addressing fundamental evolutionary dynamics, we studied the specific case of resistance to the synthetic fluoroquinolone antibiotic ciprofloxacin as an example of *de novo* resistance evolution and target modification^25,29–31^. Screening a collection of 35,211 *A. baumannii* genomes, we detected 17 distinct nucleotide variants in the quinolone resistance-determining region (QRDR)^25,32^ of *gyrA* (Fig. 1a, b). In this collection representative of the *gyrA* QRDR mutational landscape across clinical *A. baumannii* isolates (Supplementary Data 1), this is only a subset of all the theoretically possible variants: of 360 theoretical variants 261 variants code for missense changes, and of these, only 17 were observed in sequenced isolates (Fig. 1b). The most abundant allele by several orders of magnitude (Fig. 1b), was the S81L substitution, which is frequently reported to emerge under ciprofloxacin selection *in vitro* and *in vivo* ^25,30,31^ (Supplementary Table 1). We confirmed this, using a fluctuation assay with 2-8ug/mL ciprofloxacin exposure, yielding a resistance rate at the order of 1×10^−7^ (geometric mean across concentrations: 1.22×10^−7^; Supplementary Table 1). Additionally, a suite of distinct target alleles was detected between ten and hundreds of instances, as well as a few rare alleles (n<=3; Fig. 1b). While all are predicted to cluster around the conserved drug binding site of the gyrase protein complex, S81 and E85 are located most closely to the metal-ion bridge connecting to the ciprofloxacin molecule (Fig. 1a).

**Figure 1.**
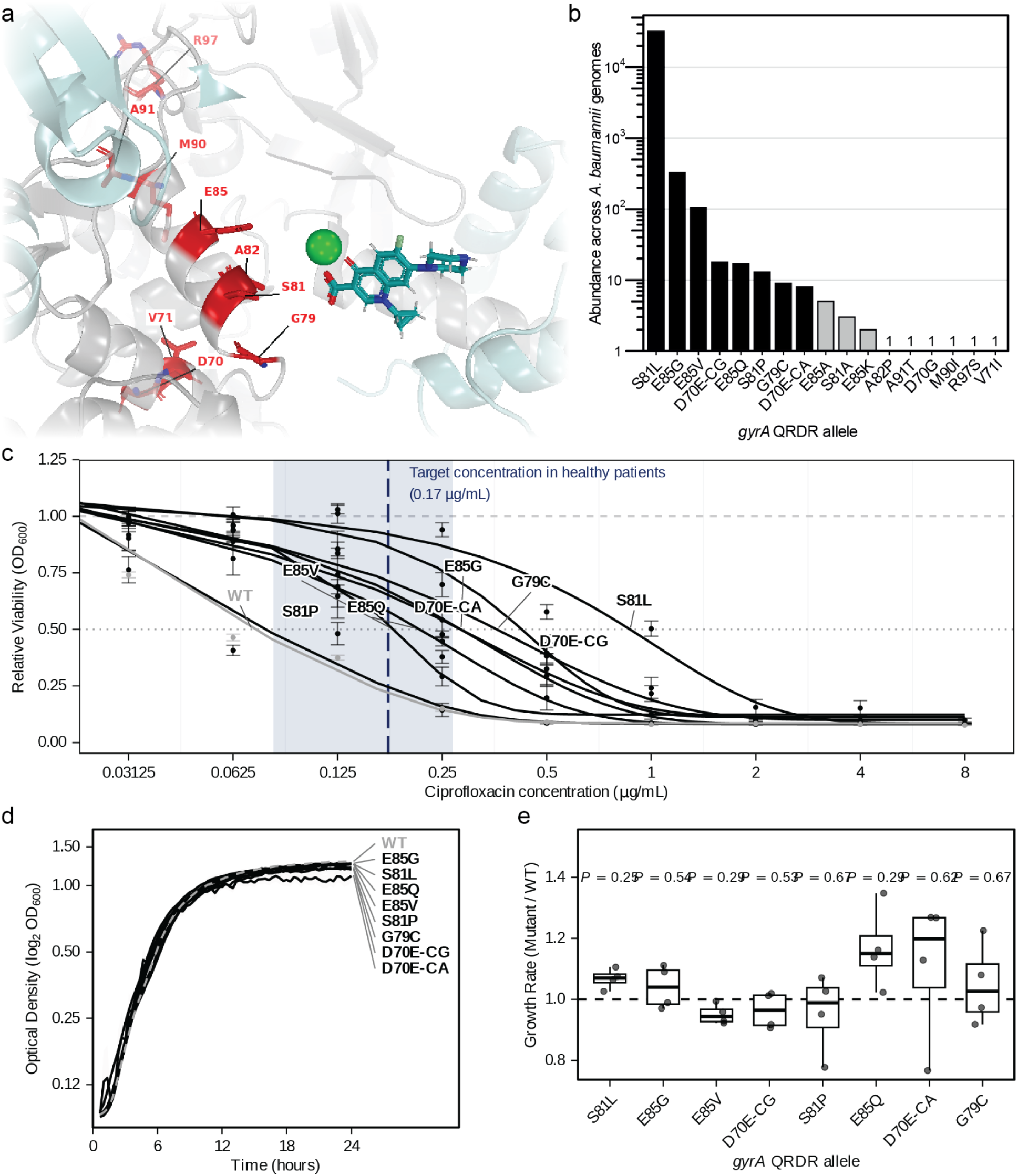
Structural, epidemiological, and phenotypic landscape of *Acinetobacter baumannii* fluoroquinolone resistance. **a**,Structural prediction of the *A. baumannii* DNA gyrase heterotetramer (GyrA₂GyrB₂) active site. The AlphaFold2-predicted complex was superimposed onto the ciprofloxacin-bound *E. coli* template (PDB: 5BTC). Mutational hotspots within the quinolone resistance-determining region (QRDR; D70, V71, G79, S81, A82, E85, M90, A91, and R97) are rendered as red sticks. A coordinating magnesium ion (green sphere) bridges ciprofloxacin (teal) to the critical S81 and E85 loci. **b,** Natural abundance of *gyrA* missense mutations identified across 35,211 *A. baumannii* genomes. Black bars denote the eight focal alleles addressed during *in vitro* characterization; grey bars indicate other observed rare alleles. D70E is explicitly split by its underlying nucleotide transversion (C>G) or transition (C>A). **c**, Ciprofloxacin dose–response curves for wild-type (WT; grey) and isogenic *gyrA* QRDR mutants (black). Data points represent mean relative viability s.e.m. (n = 4 biological replicates); solid lines show 4-parameter logistic model fits. The vertical dashed blue line and shaded ribbon indicate the simulated median target concentration (0.17 ug/mL) and population variance (0.082-0.268 ug/mL) in healthy patients dosed twice daily at 500mg ciprofloxacin orally. **d**, Drug-free growth kinetics over 24h on a log2-transformed optical density (OD600). Trajectories show mean growth curves (WT, dashed grey; mutants, solid black) with s.e.m as light grey ribbons. Allele labels are ordered by descending carrying capacity (*k*) at 24 h. **e**, Relative maximum growth rate (*r*) of *gyrA* QRDR mutants normalized to WT (dashed horizontal line). Boxplots represent median and interquartile range (IQR), with whiskers extending to 1.5xIQR; points denote individual biological replicates (n = 4). FDR-adjusted p-values above each boxplot indicate significance compared to WT in two-sided t-tests.

To test the level of resistance to ciprofloxacin conferred by >99% of the circulating *gyrA* QRDR alleles, we re-created the top 8 most frequent alleles in the susceptible *A. baumannii* lab strain ATCC 17978 using two-step allelic exchange. Log-logistic dose-response modeling confirmed a clear stratification of alleles by phenotypic resistance, with S81L displaying the profile with the greatest resistance while the S81P mutation at the same locus resulted in near wild-type susceptibility (Fig. 1c). Overlaying the *in vivo* concentration of unbound ciprofloxacin in healthy patients shows that most alleles can be detected at and above concentrations above the pharmacokinetic target of AUC24/MIC_ATCC 17978_ based on ^33,34^ (Fig. 1c).

While traditional discrete MIC testing classified both S81L and E85V as conferring an identical 8-fold resistance increase over wild-type (Supplementary Data 3a), continuous log-logistic dose-response modeling revealed that E85V exhibits a lower IC50 than S81L (Fig. 1c). This divergence highlights the limitation of single-point MIC measures and confirms that continuous IC50 kinetics provide a more nuanced measure of concentration-dependent selection under dynamic drug exposure. Additionally, the two reconstructed D70E mutants showed distinct dose-response profiles at sub-inhibitory concentrations of ciprofloxacin (<1*μ*g/mL ciprofloxacin), although genomically identical beyond the introduced synonymous mutations (Fig. 1c, Supplementary Data 2, Supplementary Table 2). Complementing the dose-response curves with the clinical microbiology approach of scoring minimum inhibitory concentrations (MICs) by eye revealed that 5 of the 8 reconstructed alleles would be considered strongly resistant in comparison to the ATCC 17978 WT (MIC ≥ 4x WT MIC; Supplementary Data 3) and that bactericidal effect of the drug was maintained (Supplementary Data 3). Despite the clear differences in resistance, we did not detect any major costs in Malthusian fitness, i.e. bacterial growth, as there were no significant differences in growth rates in the absence of ciprofloxacin between the resistant variants and the susceptible wild type (Fig. 1d,e). In the absence of drug pressure, different resistance alleles would thus be expected to co-exist and despite S81L showing the greatest resistance of all alleles, it might be outcompeted by E85Q in the absence of drug pressure, as this allele reached the the maximal growth rate in drug-free medium (Fig. 1e). Taken together, this highlights that there are multiple target modifications of *gyrA* that lead to clinically-relevant levels of resistance, yet no significant fitness cost under the defined growth conditions. While the level of resistance will determine survival under antibiotic exposure, a mutation has to occur in the first place. Given that not all mutations are equally likely to occur due to biochemical and evolutionary constraints^17,18,35^, they may easily outcompete those with the highest potential resistance^5,14,36–38^. To build towards a *de novo* predictive framework of the emergence and success of antibiotic resistance alleles, we therefore performed a retrospective analysis of *A. baumannii* evolution to assess and quantify the impact of mutational bias on *gyrA* evolution in this pathogen.

### Multiple evolutionary drivers historically explain allelic abundance

To build a model for the evolutionary success of resistance alleles, we first validated our understanding of the drivers of resistance with a retrospective evolutionary analysis. Specifically, we aimed to integrate the likelihood of resistance variants occurring and then surviving under treatment. As the chance of a resistance mutation appearing is determined at the level of the genome, we first inferred mutational biases in *A. baumannii*. For this, we chose a phylogenetically-informed approach that considers the nucleotide changes in independent mutational events rather than spread of mutations during transmission^37^. We reconstructed the evolutionary history of *gyrA* across a core-genome phylogeny based on 2,443 high-quality isolates to map independent parallel mutations against neutral genomic substitution biases and confirmed that the studied *gyrA* variants are spread across the species tree (Fig. 2a). Ancestral state reconstruction showed strong evolutionary convergence, with the primary S81L driver arising independently over 100 times and rarer variants (e.g., E85Q, G79C, S81P) appearing almost exclusively as secondary modifications in S81L background (Supplementary Data 4). However, our overall approach assumes that drug developers can identify individual mutations that lead to resistance, while they have no a priori knowledge on their potential sequence of emergence. Consequently, we continued treating the *gyrA* variants as equivalent as would be the case at the stage of comparing single mutants during drug development. To assess whether common variants would be favored already at the genomic level, we evaluated whether mutational bias inherent to the *A. baumannii* genome as a predictor of the abundance of specific *gyrA* QRDR mutant alleles (Fig. 2b). Regression analysis confirmed that substitution likelihood relates significantly with the abundance of a resistance mutation in this sequence stretch (log10-transformed counts: n = 14, R^2^ = 0.37, F_1, 12_ = 7.09, P = 0.021). This links the availability and biochemical access to make any of the possible nucleotide exchanges to their likelihood of emerging. This predictive relationship between substitution likelihood and allele abundance is specific to the quinolone resistance-determining region (QRDR), where mutations confer selective advantages under drug pressure (Supplementary Table 3). Outside the QRDR the association is absent rather than merely diluted (R² = 0.003, P = 0.43), and it does not hold across the full *gyrA* coding sequence (R² = 0.007, P = 0.22; Supplementary Table 3), confirming that mutational bias likely acts in tandem with target-site drug selection.

**Figure 2.**
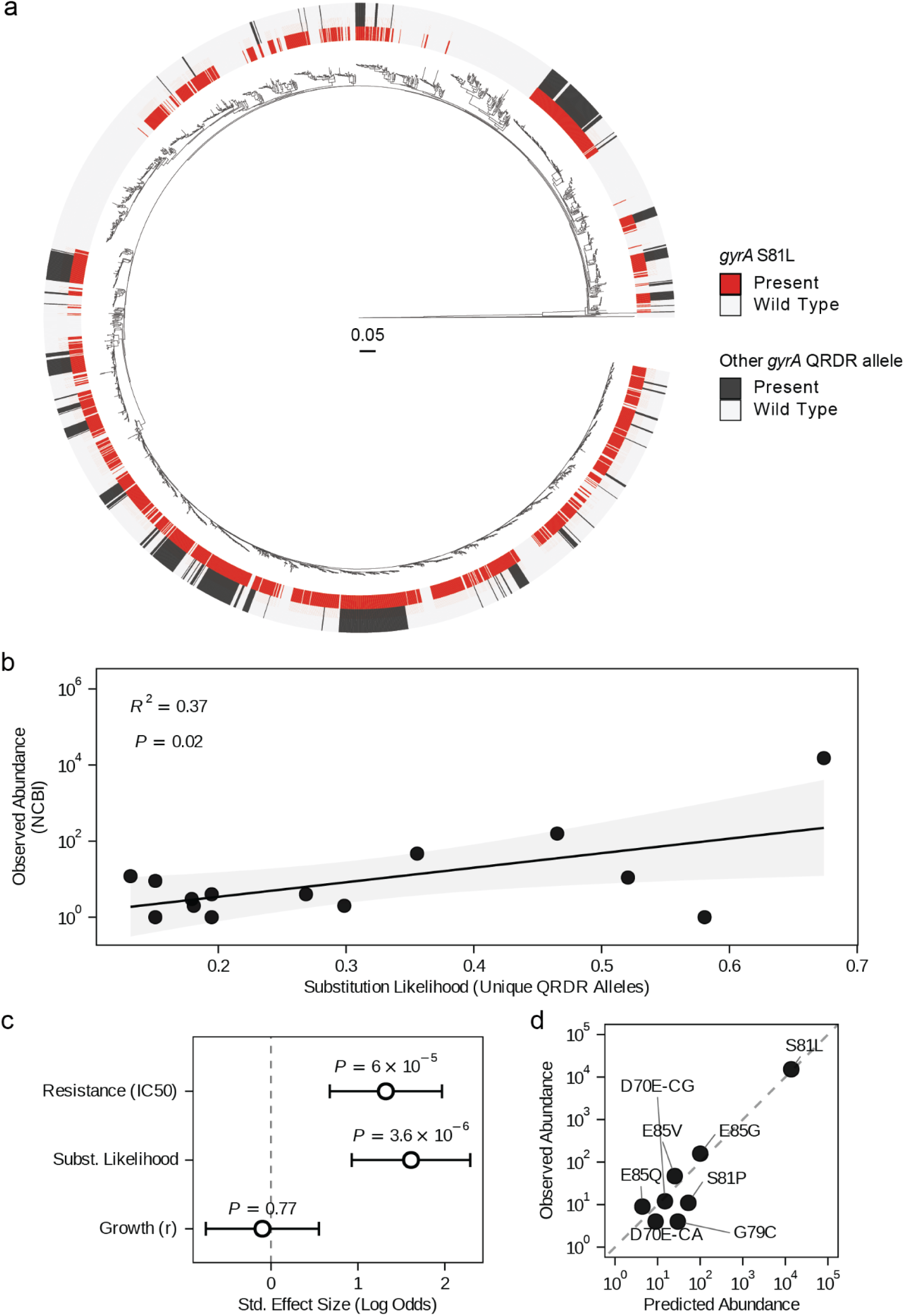
Phylogenetic context and mechanistic predictors of *Acinetobacter baumannii gyrA* QRDR variant evolution. **a**,Core genome maximum-likelihood phylogeny of 2,443 high-quality *A. baumannii* genomes, rooted to the outgroup *A. baylyi* ADP1. The inner ring designates the presence/ absence of the most common *gyrA* QRDR mutation S81L (in red vs. wild type in light grey). The outer ring displays presence/absence of other *gyrA* QRDR alleles (in dark grey vs. wild type in light grey). The scale bar represents nucleotide substitutions per site. **b**, Linear regression with 95% confidence interval between global genomic abundance (log10-count of NCBI *A. baumannii* genomes and substitution likelihood across unique *gyrA* QRDR missense unique nucleotide substitutions (n = 14, R^2^ = 0.37, F_1, 12_ = 7.09, P = 0.021)). **c**, Standardized effect sizes (log-abundance) from the top-performing multivariate Negative Binomial generalized linear model (GLM) predicting allele abundance. Error bars show 95% confidence intervals. *P* values were inferred using two-sided Wald tests. **d**, Model validation of predicted vs. observed genomic abundance by the GLM with a dashed unity line.

In contrast to other genes commonly observed to mutate under antibiotic selection, *gyrA* showed a particularly strong response to mutational bias (Supplementary Data 5). Profiling variants in previously identified driver mutations in *pmrB* and *adeS*, linked to colistin and tigecycline resistance respectively^25,30,39–41^, showed no significant impact of substitution likelihood on the abundance of specific resistance alleles across the 35,211 *A. baumannii* genomes from NCBI (Supplementary Data 5). When *A. baumannii* evolved under controlled lab conditions in a morbidostat bioreactor, however, substitution likelihood significantly explained the abundance of *pmrB* variants. For *gyrA*, only a single *gyrA* variant, the mutationally likely S81L, was detected, while *adeS* variation showed no relationship to mutation bias (Supplementary Data 5).

When comparing laboratory evolution to the global genomic surveillance of *gyrA* QRDR variants (Fig. 1b), we note that the NCBI genome collection(n = 35,211) represents a sampling depth several orders of magnitude greater than the isolates sampled from the laboratory morbidostat bioreactors^30,41,42^, thus not expected to exhaustively sample all 17 rare QRDR target variants observed across global clinical collections. These data highlight that mutational bias impacts mutant success differently in *in vitro* bioreactor conditions versus evolution in a host with fluctuating drug concentrations. Integrating emergence and selection of *gyrA* QRDR resistance alleles, we assessed the impact of substitution likelihood, resistance, as well as growth cost on resistance allele abundance. A multivariate Negative Binomial Generalized Linear Model (GLM) integrating all three biological constraints successfully predicted global clinical frequencies (P < 0.001, Fig. 2c, d). We confirmed the statistical independence of these predictors through Variance Inflation Factor analysis (VIF_IC50_ = 1.24, VIF_subLikelihood_ = 1.30, VIF*_r_* = 1.09), verifying that resistance levels and mutation biases act as synergistic, uncoupled constraints.

### Stochastic multi-scale modeling captures dynamic selective filters and predicts priority resistance variant

To transition from retrospective correlations to the potential for prospective forecasting, we integrated our biophysical and phenotypic measurements into a continuous-time stochastic modeling framework that simulates evolutionary trajectories under real-world pharmacokinetic fluctuations for fluoroquinolones (Fig. 3a). In a nutshell, we simulated the emergence and selection of *gyrA* QRDR variation in *A. baumannii* infection across a patient population treated with ciprofloxacin. In this, we parameterized the emergence probability and resistance level of the tested *gyrA* QRDR based on the empirical data collected above. Given the lack of significant differences in growth between the tested resistance mutants (Fig. 1d,e), we assumed uniform growth parameters in the absence of drug treatment. Including dynamic antibiotic concentrations in the model not only mirrored patient physiology, but allowed for more permissive selection windows, and improved the model fit as compared to static antibiotic exposure (Supplementary Data 6). Comparison of modeling architectures revealed that implementing dynamic, instantaneous kill rates (α(*t*) ), provided a significantly superior fit compared to static, time-averaged implementation, where only the dynamic model successfully captured the “sawtooth” population dynamics of mutant expansion during PK troughs where drug concentrations fall below the mutant MIC (Supplementary Data 6). Seeding individual patients with an initial burden of 50 bacterial cells, we allowed infected bacterial populations to grow until reaching a bacterial load of 10^7^ cells, then initiating antibiotic treatment. We defined a clinical rebound based on three concurrent criteria at day 14: (1) a 2-log10 expansion from the post-treatment population nadir^20,43^ (or reaching a peak ≥ 10^2^ cells if the nadir was zero), (2) a peak mutant burden ≥ 10^4^ cells, and (3) a final total surviving population ≥ 10^3^ cells (see materials and methods). To evaluate how parameter changes affect model predictions and optimize parameter fit, we performed a one-at-a time sensitivity analysis. To avoid tuning parameters to empirically observed allele distributions, we calculated how parameter changes (in initial pathogen load, kill rate scalar ɑ, mutation rate *μ*, growth rate scalar *r* or the turnover rate *T*) would affect treatment efficacy, i.e. the rate of rebounds (Supplementary Data 7). While the empirically-informed initial parameter set led to nearly complete treatment efficacy across patients, decreasing the kill rate scalar led to realistic rebound rates between 1-5% in healthy patient cohorts^44,45^ (Fig. 3b, Supplementary Data 7a, b). We therefore proceeded in modeling with an intermediate kill rate scalar of 0.25, likely representing the translation of *in vitro* killing activity to *in vivo* efficacy of the drug and implemented the simulations starting with the complete set of reconstructed, viable *gyrA* QRDR alleles (Fig. 3a).

**Figure 3.**
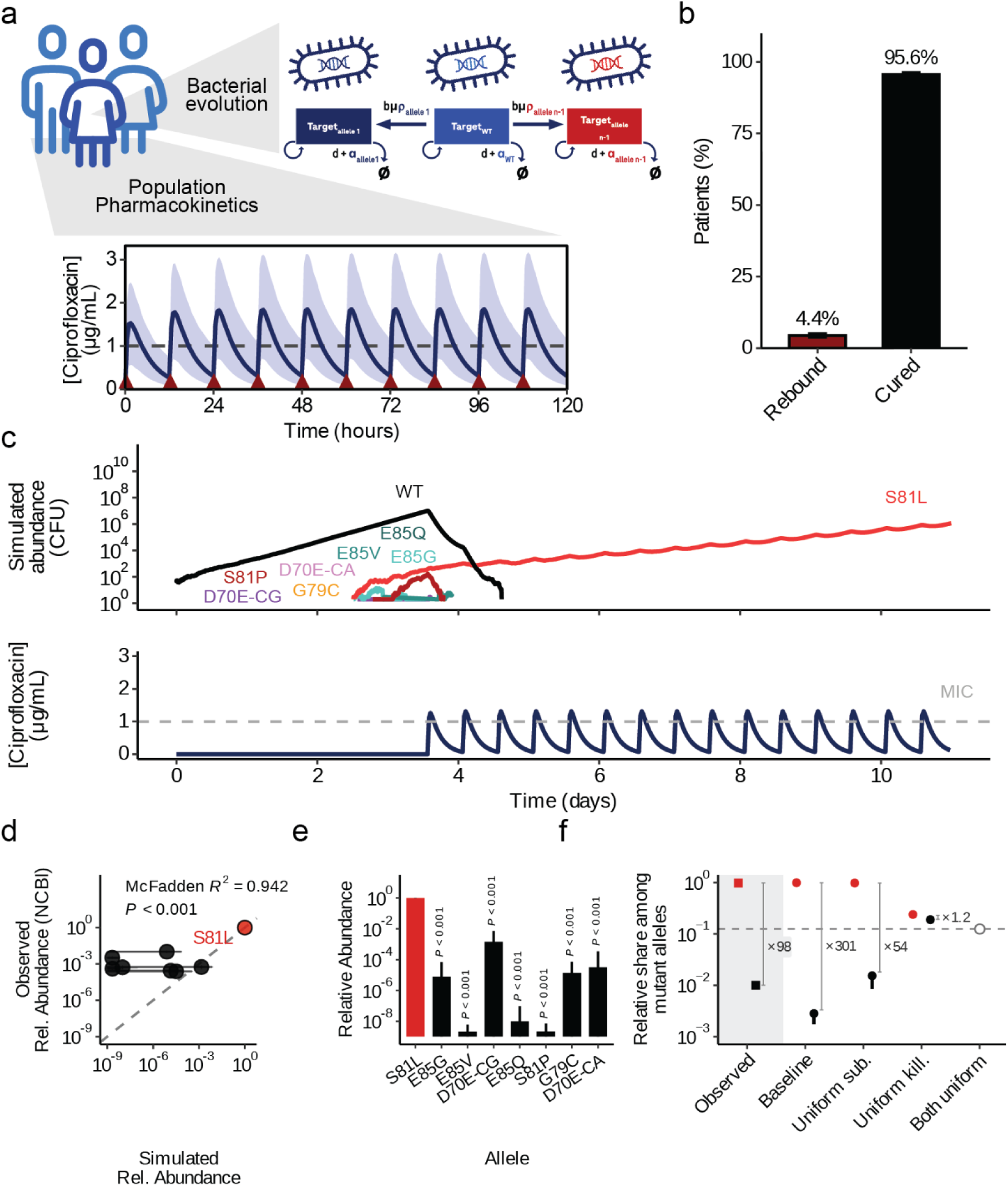
Mechanistic multi-scale modeling integrates pharmacokinetics and evolutionary dynamics to predict clinical AMR prevalence. **a**,Schematic overview of the modeling framework. Virtual patient populations undergo pharmacokinetic (PK) simulations of standard ciprofloxacin dosing (bottom inset; dark blue line represents the median concentration over time in healthy patients, the shaded region the population variance, red triangles indicate twice daily dosing at 500 mg). The dynamic drug concentration, *C*(*t*), informs a stochastic birth–death evolutionary model (top inset), where wild-type (WT) and mutant *Acinetobacter baumannii* alleles compete based on intrinsic replication (*b*), natural death (*d*), global mutation rate (*µ*), allele-specific substitution likelihood (*p*), and drug-induced killing (*α*). **b,** Simulated clinical outcome frequencies for the healthy baseline cohort (*N* = 100,000 virtual patients). Bar heights indicate the median percentage of patients classified as rebound or cured across simulation batches; error bars denote ±1 s.d. **c,** Representative *in silico* virtual patient trajectory with relapse. The top plot displays the stochastic population dynamics of the WT (black) and emerging *gyrA* QRDR variants (colored lines). The bottom plot shows the corresponding individual ciprofloxacin concentration profile relative to the wild-type ATCC 17978 minimum inhibitory concentration (dashed line = MIC). **d**, Concordance between simulated relative allele abundance in the post-treatment resistant reservoir and observed global prevalence from *A. baumannii* NCBI genomes (N = 32,545 isolate allele observations). The red point represents S81L, while all other non-WT gyrA QRDR alleles are shown in black; the dashed line indicates 1:1 identity. Goodness-of-fit is evaluated via multinomial log-likelihood estimation against a uniform null model (McFadden’s R^2^ and Likelihood Ratio Test P-value). Horizontal error bars denote inter-patient population dispersion (+/- 1 s.d.) across simulation batches; vertical error bars denote empirical genomic sampling uncertainty (25th - 75th percentile interquartile range across 1,000 bootstrap replicates). **e,** Predicted relative abundance of each *gyrA* QRDR allele in the resistant reservoir under the baseline model (log10-scale). Error bars denote inter-patient population dispersion (+/- 1 s.d.) across simulation batches. Two-sided Mann–Whitney U tests compared S81L relative abundance against each alternative allele and resulting by FDR-corrected P-values are shown. **f,** Model component attribution by parameter ablation. For the baseline model and for each ablated scenario, points show the share of the simulated post-treatment resistant reservoir held by S81L (red) and by the most abundant non-S81L allele (black, runner-up depending on the simulation), alongside the corresponding observed shares among NCBI genomes (filled squares, shaded panel at left). Allele-specific drug-induced kill rates (“Uniform kill.”, set to α_S81L_) or allele-specific nucleotide substitution likelihoods (“Uniform sub.”) are replaced by a single value shared across alleles. Error bars denote +/-1 s.d. across five independent replicate simulations of 100,000 virtual patients each. Grey brackets and annotations give the dominance margin between S81L and the runner-up allele, defined conservatively as the minimum S81L share across replicates divided by the maximum share attained by any non-S81L allele in any replicate. The dashed horizontal line and open circle mark the chance expectation 1/K = 0.125 for K = 8 exchangeable alleles, i.e. uniform allele proportions.

Tracking individual *in silico* patient trajectories exposed the complex, time-resolved population bottlenecks that govern variant establishment and success. Following drug administration, the wild-type population rapidly crashed, opening a critical selective window with pharmacokinetic troughs where concentration dropped below variant-specific MICs (Fig. 3c). Simulating the baseline healthy cohort, this led to an overall high treatment success with a realistic rebound rate of 4.4% (Fig. 3b). Focusing on these rebounding patients, we analyzed the total resistance reservoir by calculating the Area Under the Curve (AUC) of the prevailing resistance alleles across the observed treatment window. Without explicit parameter fitting to historical clinical frequency counts, simulations across 100,000 virtual patients showed that S81L outcompetes all alternative alleles by multiple orders of magnitude (Fig. 3d, e). A complementary literature synthesis across preclinical and clinical studies confirmed that our framework allowed prioritizing the most abundant clinical variant prospectively, specifically identifying the winning allele’s competitive advantage (Fig. 1b, Supplementary Table 1). In contrast, the relative abundance of the more rare *gyrA* QRDR alleles could not be predicted, given the large standard deviations across patients (Fig. 3d, e). Since rarer QRDR variants occur in clinical genomes almost exclusively as secondary modifications of an S81L background (Supplementary Data 4), a process this first-step model deliberately excludes, we restrict our quantitative claims to the identity and the dominance of the winning allele. Formally ranking the non-S81L mutant alleles is not possible in this case, but we do note that the model predicts their occurrence (Fig. 1c, Fig. 3e). To decouple the influence of the individual biological variables that generate this landscape, we conducted a model component attribution analysis in which each allele-specific parameter was replaced by a uniform value across alleles (Fig. 3f, Supplementary Data 8, 9). While S81L remained the most abundant allele across scenarios, replacing the allele-specific drug-induced kill rates with a single uniform value (set to α_S81L_) collapsed the predicted dominance of S81L. The margin between S81L and the next most abundant allele fell from ×301 in the baseline model to ×1.2, against ×98 observed among NCBI genomes, i.e. to within a factor of 1.2 of the chance expectation (i.e. 1/K for K = 8 exchangeable alleles; Fig. 3f). Ablating the allele-specific nucleotide substitution likelihood instead left S81L dominant, with a margin of ×54. Across five independent replicate simulations S81L remained the most abundant allele in every replicate of every simulated scenario, including both single-parameter ablations (5/5 in each). Thus, either resistance or nucleotide substitution likelihood alone are sufficient to identify the winning allele and neither is individually necessary. The same ordering is recovered on the multinomial log-likelihood scale, where uniform kill rates degrade the per-isolate fit by 1.23 log-likelihood units (replicate range −1.28 to −1.18; worse than baseline in 5 of 5 replicates and in 1,000 of 1,000 genome resamples), and is robust to clonal outbreak overrepresentation and temporal sampling bias when the analysis is bootstrapped over 1,000 iterations stratified by Pasteur sequence type and collection era (Supplementary Data 8). Replacing allele-specific parameter values with uniform mutant drug-specific kill rates (set to *ɑ_S81L_*) degraded the multinomial model fit by 1.21 log-likelihood units per isolate (95 % bootstrap interval -1.23 to -1.20), with the ablated model fitting worse than the baseline in 1,000 of 1,000 resamples. We confirmed this was robust against clonal outbreak overrepresentation and temporal sampling biases in public repositories by bootstrapping the ablation analysis over 1,000 iterations stratified by Pasteur sequence type and collection era (Supplementary Data 8).

Using the benefit of hindsight, we repeated the simulations with a focus on *gyrA* QRDR alleles known to evolve as first mutations, i.e. primary alleles (Supplementary Data 4 and 9). Here too S81L was identified as the winning resistance allele, outcompeting the remaining primary alleles by ×2,692 in the baseline model against ×98 observed, and remaining the most abundant allele in 5 of 5 replicates in every simulated scenario. The ablation analysis again identified the allele-specific kill rate as the dominant contributor, reducing the S81L margin to ×1.35, while the contribution of the allele-specific substitution likelihood again could not be resolved above run-to-run variation (Supplementary Data 8 and 9).

While this baseline model captures the average evolutionary trajectory in a healthy, standardized cohort, the impact of individual physiological variability remained unknown. We therefore extended this framework to investigate how variations in pharmacokinetic profiles, treatment adherence, and host immunity drive resistance emergence and landscape across distinct patient populations.

### Highly abundant winner allele can be identified across host physiologies and treatment adherence

While the baseline model successfully captured average evolutionary trajectories, it assumes a standardized healthy host environment that rarely reflects clinical reality. To move beyond this average, we extended our framework to investigate how individual physiological variability, host immune status, and treatment adherence drive resistance emergence across distinct patient populations (Fig. 4a). In the clinic, these simulated cohorts mirror patient physiologies in intensive care units, where *A. baumannii* predominantly manifests as pneumonia^46^, with variable respiratory drug penetration, alongside common kidney impairment^47^ or secondary immunosuppression^47–49^, leaving uncompromised baseline hosts as a small minority. We utilized a “virtual twin” strategy, applying specific physiological modifiers such as altered drug elimination (k_el_), absorption (k_a_), or immune clearance (δ), to mirror populations contrasted with the baseline healthy population. Simulated patients experiencing low drug penetration (i.e. simulating restricted ciprofloxacin access to the site of infection) suffered a strongly increased 52.6% clinical rebound frequency compared to healthy patients (Fig. 4c). In these individuals, the reduced drug concentrations created a sustained, highly permissive selective window that allowed more bacteria and thus multiple alleles to remain (Fig. 4c). Conversely, simulated renal impairment, under which drug exposure increases if doses are left unadjusted, resulted in near universal cure rates, thus limiting the resistance reservoir (Fig. 4b-d). Notably, modeling a single skipped evening dose led to a clinical rebound frequency of 5.98% (Fig. 4c). While there are many variations of non-compliance, we deliberately fixed the skipped dose at the second dose (the first evening dose), as these might be the most prone to being missed in routine variability^50^. As a consequence, simulating a skipped dose this early in the regimen allowed us to evaluate the evolutionary consequences of a drug trough during peak bacterial burden and high mutational supply. Tracking these specific *in silico* trajectories confirmed that even a transient, single-dosing sub-therapeutic window was sufficient to rescue clearing bacterial populations and catalyze a trajectory toward relapse (Fig. 4b). As expected from clinical observations^44,51^, neutropenic virtual patients showed a moderate increase in rebound rate compared to healthy patients (Fig. 4c) and acted as multi-allelic reservoirs including rare resistance alleles (Fig. 4d; ^31^). Comparing bacterial burdens across simulated patient groups consolidated that neutropenic and low-penetration cohorts contribute the largest total resistant biomass (Fig. 4b). Therefore, these patients are expected to significantly elevate the risk of environmental shedding and transmission compared to healthy cohorts.

**Figure 4.**
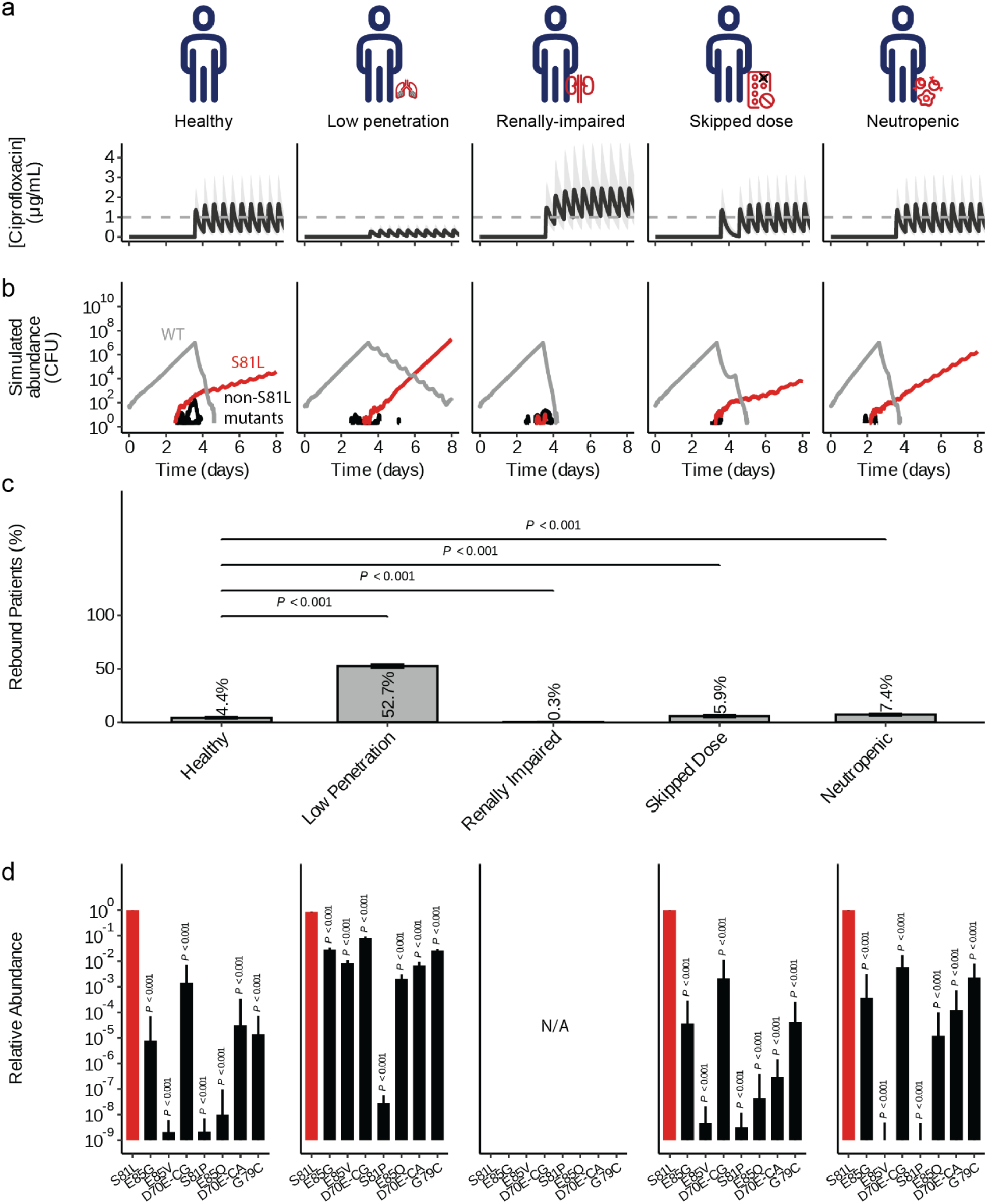
Host physiological variability, immune competence, and treatment non-adherence alter macro-clinical outcomes and break target-site allelic landscapes. **a**,Pharmacokinetic (PK) profiles across the simulated virtual patient cohorts (N = 100,000 virtual patients/ group). Black lines represent the median ciprofloxacin concentration (ug/mL) over time at the site of infection with population variance shown in grey. Dashed gray lines mark the MIC of WT susceptible ATCC 17978. **b,** Representative *in silico* time-resolved population trajectories (CFU) over an 8-day timeline for each distinct cohort. The 8-day window was selected to illustrate the timeframe of peak population collapse, mutant selection, and early dynamic rebound, whereas cohort-wide clinical outcome frequencies in c were evaluated at day 14 as in Fig. 3). Grey lines track the rapid clearance or escape of the wild-type population; red and black lines track the stochastic emergence, extinction, or fixation of individual *gyrA* QRDR resistance variants. **c,** Population-wide clinical rebound frequencies (%) across the five virtual cohorts at day 14. linical rebound is defined by three concurrent criteria: a ≥ 2-log10 expansion from the post-treatment nadir (or peak ≥ 10^2^ cells if nadir is zero), a peak mutant burden ≥ 10^4^ cells, and a final total population ≥ 10^3^ cells. Error bars denote 1 s.d. across simulation batches; P-values are derived from pairwise two-sided chi-squared tests relative to the healthy baseline cohort. **d,** Relative abundance profiles of surviving *gyrA* QRDR variants within the post-treatment resistant reservoir across the rebounding cohorts. The primary driver allele S81L is highlighted in red, alternative variants in black. Error bars indicate 1 s.e.m. across simulation runs. The renally impaired cohort is designated as Not Applicable (N/A) due to a negligible rebound rate. The emergence of distinct signatures in the neutropenic panel visually confirms the rescue and survival of low-fitness, high-resistance variants following the removal of host immune filters.

Across the simulated patient groups and exposures, our model faithfully recovered the clinically significantly most successful allele (Fig. 4d). This aligns well with the dominance of S81L across real-world sampled genomes (Fig. 1a, b). Collectively, these findings demonstrate that host physiology and ecology can shape the specific structural landscape of clinical failure, while genetic accessibility and selection drive the success of the winning allele. While we here implemented simulations with *A. baumannii* and ciprofloxacin-specific empirical parameters, the model itself presents a general framework that can be adapted to other bacteria and antibiotics with respective resistance, growth and PK data. This multi-scale framework thus provides a robust quantitative platform to stress-test and de-risk emerging antibiotic candidates across heterogeneous patient populations prior to entering clinical evaluation.

## Discussion

This study establishes a quantitative, multi-scale framework that links within-host selection dynamics, mutational accessibility, and patient physiology to forecast the evolutionary success of antibiotic resistance variants. Rather than treating sweeping resistance as an emergent property to be observed after the fact, our results show that successful resistance alleles can be anticipated from measurable biological constraints. To leverage evolutionary predictability when (i) the mutational target size is restricted, (ii) the mutational supply is limited, and (iii) selection pressures are well-characterized across time^8,52^, we established a forecasting model to prioritize resistance alleles that may appear in the clinic. Focusing on fluoroquinolone resistance in *A. baumannii* as a proof-of-concept system, we identified three interacting constraints that determine evolutionary trajectories across scales of biological organization. These include mutational accessibility at the nucleotide level, allele-specific fitness under dynamic drug exposure, and ecological filtering imposed by host immunity. The success of the clinically most wide-spread *gyrA* S81L mutation can be explained by the aligned high probability to emerge as well as the selective advantage this allele has over other viable target variants. More generally, comparing resistance-mediating genes *gyrA*, *pmrB* and *adeS* suggested that selective filters are overwhelmed by mutational biases to different extents in across resistance-mediating genes. In the following, we will discuss (i) the role of mutational bias and selection in shaping bacterial AMR evolution, (ii) the predictive boundaries of our modeling workflow and (iii) the impact of PK shifts on outcomes across patients. Thus, demonstrating how multi-scale modeling from first principles has potential to prioritize the clinical relevance of future resistance alleles.

Our analysis demonstrates that substitution likelihood defines which resistance alleles will emerge frequently and can thus directly affect resistance evolution^14,17,36,37^. In the case of antibiotic target gene *gyrA*, the most commonly observed variant both *in vitro* as well as the clinic, S81L, is reached by the most likely nucleotide substitution in the *A. baumannii* genome, a C to T transition. While the resulting resistance change alone makes this allele more competitive under drug exposure, we statistically confirmed that the mutational bias independently contributes to the success of this allele. Furthermore, we corrected for parallel evolution rather than biases stemming from the spread of individual clonal outbreaks by phylogenetically correcting for relatedness of isolates in our statistical and modeling analyses. This underscores the two-step process of evolution: variants first have to emerge, which is proportional to their respective nucleotide substitution likelihood, and then be selected, in this case survive antibiotic treatment. As our framework simulates exposure to a novel drug, it starts with a susceptible, wild type population of pathogens that evolve resistance *de novo* and are then exposed to a bactericidal antibiotic. Consequently, our framework relates to early infection stages, or under strong pharmacokinetic bottlenecks, where population sizes are reduced and stochastic establishment dominates^21^. Importantly, differences in resistance between two nucleotide variants leading to the same amino acid change, like for *gyrA* D70E, highlighted that forecasting needs to explicitly consider synonymous variation and not only structural changes that affect drug interactions at the protein level^16,53^. While the mechanistic basis remains unclear, but could be linked to codon bias, secondary mRNA structures, translation kinetics amongst others^54,55^, it reinforces that predictive models must incorporate nucleotide substitution biases rather than assuming uniform accessibility of amino acid space. Similar to observations in resistance development in other kingdoms of life^36^, this shows that those mutations that are likely to appear early and often and at the same time provide sufficient resistance have the capacity to spread most effectively, e.g. in *M. tuberculosis* antibiotic resistance^37^, kinase inhibitor resistance in ABL1+ leukemic cancer^5^, or HIV reverse transcriptase mutations^56^. In comparison to more slowly replicating cancer cells, such a significant influence of mutational bias on the resistance landscape is less likely, given that rapidly dividing bacterial populations might enrich resistance alleles more quickly^5^. This might indicate that nucleotide substitution likelihood is potentially a more general pre-selective filter across human pathogens.

While our modeling leverages the data collected during the antibiotic development process, its translational limitations also affect its predictive power. Firstly, we assumed that during development of a novel antibiotic, there is no *a priori* knowledge about the sequence of potential, viable resistance mutations before bacteria are exposed. Therefore, we did not filter any of the top most frequent *gyrA* QRDR variants observed to be viable in the real-world and reconstructed them in the isogenic ATCC 17978 background. While this allows us to directly focus on the specific, chemically-addressable mutations and mirrors mutant identification and validation during resistance screening, it purposefully simplifies the evolutionary process surrounding resistance. Considering the initial *gyrA* substitution as the rate-limiting evolutionary step towards resistance is biologically justified by the physiological state transition associated with first-step resistance demonstrated in *A. baumannii*^26^. Our model therefore focuses on *gyrA* QRDR mutations and specifically excludes second step mutations in other genes, e.g. mutations in *parC* commonly observed in tandem with changes in *gyrA*^26,57^. Distinguishing between primary and secondary mutations within the same gene confirmed resistance as the known key driver of allele success, but showed that genome-level nucleotide substitution likelihood further refined forecasting primary allele success. Without the benefit of hindsight, we thus conclude that identification of spontaneous mutations or genetic screening should be integrated with observations on mutational sequence during serial passaging to identify potential contingencies. Secondly, we established a general evolutionary framework based on a generalizable development workflow, yet the additional predictive advantage of mutational bias appears resistance-gene-specific. While mutational bias independently drove allele success at the target locus *gyrA*, it did not correlate with SNP abundance in *pmrB* or *adeS*. This difference illustrates that mutational supply is likely most decisive for active-site target modifications where mutational target space is highly constrained^14,17^. While the three loci in focus here, i.e. *gyrA*, *pmrB*, and *adeS* illustrate a range of influence of mutational bias, future studies might expand the platform to additional direct target pairs (e.g., *rpoB*). Thirdly, we focused on fundamental evolutionary and physiological drivers of resistance allele success in an infection-agnostic model. Drawing from *A. baumannii* genomes across infection types and body sites as well as pharmacokinetic parameters based on meta-analyses of clinical data^33^ allowed us to generally validate our model forecasts against the broad landscape of ciprofloxacin and fluoroquinolone resistance. Patient cohort simulations, however, already hint at the potential of fine tuning the modeling to specific target populations and infection types. Together, the predictive power of our proof-of-concept for the direct drug-target interactions governing GyrA and fluoroquinolones, is defined by clear boundary conditions and offers a quantitative framework that can now be applied and tested across other direct target-drug pairs and infection contexts. Depending on the antibiotic candidate molecule, structural changes in the drug target are chemically addressable by rational drug design^5,58^, thus making it immediately relevant for next-generation antibiotic design. However, selective filtering of antibiotics, as well as the host environment, may shape or overwhelm the impact of resistance and mutational bias on the finally observed resistance landscape.

For resistance forecasting from the lab to patients, physiological measures need to be integrated with bacterial evolutionary dynamics. Standard PK/PD metrics (e.g., PK drivers like AUC/MIC) obscure the temporal structure of selection and impact of specific host physiology^59^. Troughs in antibiotic concentration, for example, define the mutant selection window, where resistant subpopulations can expand while susceptible populations are suppressed^20,51^. Due to the rapid bactericidal effect of ciprofloxacin, rebounding resistance alleles reflect the mutational supply already present in the large pre-treatment bacterial load before the first dose is administered.

Within this framework, within-host ecology acts not as a dynamic generator of diversity, but as a severe purifying filter layered on top of this pre-existing stochastic mutant reservoir, where human pharmacokinetic fluctuations and immune clearance determine whether a baseline variant survives the initial crash. We confirmed that reduced drug penetration increases failure rates and expands resistance reservoirs, and neutropenic conditions relax ecological filtering, allowing otherwise unfit variants to persist. While we modeled immune impairment and reduced drug penetration by lowering drug exposure or the bacterial kill rates in our model, physiological PK/PD modeling may serve as a role model to include non-linear modulation of exposure or dynamic immune feedback during infections^44,60,61^. Already now, the framework presents both a testing ground to compare efficacy as well as quality of resistance control across patient groups, and thus inversely a potential guide to resistance-conscious dosing across patient groups.

Together, our findings demonstrate that resistance evolution is a property of the host–pathogen–drug system, rather than any individual component^62^. While mutation rates alone are insufficient to predict resistance risk, we showed that mutation bias, fitness, and dynamic selection pressures must be integrated to identify high-risk variants. This framework supports a transition toward model-informed drug development for resistance forecasting and de-risking of antibiotics^4^. While conventional preclinical workflows rely on static mutation frequency assays, candidate antibiotics frequently suffer rapid resistance emergence across ESKAPE pathogens due to standing genetic variation and uncharacterized mutational supply^63^. We therefore present a resistance-conscious drug development workflow with the following five steps: (i) *in silico* inference of nucleotide substitution likelihoods based on sequenced genomes, followed by (ii) identification of viable alleles in resistance genes (e.g. by classic spontaneous mutagenesis, targeted mutagenesis or even systematic deep mutational scanning^58,64–66^), and (iii) *in vitro* phenotypic and growth kinetic profiling (IC50, growth rate *r*) of isogenic mutants, (iv) collection of PK parameters specific to the antibiotic, either from early phase clinical studies or dynamic models like the hollow-fiber infection model^67^, and finally (v) stochastic simulation of bacterial evolution across patient cohorts to prioritize must-hit resistance alleles. Applying this workflow should help identify likely evolutionary trajectories, and thus help guide what resistances to target with next-generation molecules or which ones to deprioritize during rational drug design. As a result, this model-driven forecasting should actively shorten the feedback loop of clinical relevance as it may prioritize future resistance alleles before active clinical use of a novel antibiotic. Moreover, varying dosing regimens in the underlying PK simulations can be used to optimize dosing regimens minimizing resistance across patient groups. Inversely, by varying the patient population size studied, the tool may be used to investigate the minimal clinical trial size to confidently infer future resistance signatures. Taken together, multi-scale evolutionary modeling thus provides a quantitative basis for integrating resistance forecasting into clinical decision-making.

## Materials and methods

### Structural modeling and visualization

To map the spatial distribution of mutational hotspots within the *A. baumannii* ATCC 17978 target complex, the tetrameric GyrA-GyrB structure was predicted using ColabFold^68^ v.1.5.5. The highest-ranked model was structurally superimposed onto the high-resolution reference crystal structure of the *Escherichia coli* DNA gyrase complexed with ciprofloxacin and DNA (PDB: 5BTC) using PyMOL^69^ v3.1.6.1. Spatial filtering was applied to isolate the quinolone resistance-determining region (QRDR); specifically, residues D70, V71, G79, S81, A82, E85, M90, A91, and R97 (identified using variant calling across *A. baumannii* genomes described below) were rendered as sticks. A 10 Å radial proximity filter centered on the primary resistance driver, Ser81, and a 5 Å filter anchored to the ciprofloxacin molecule were utilized to map the mutational search space. The final structural orientation was locked using a defined 20 Å sphere focal view and residues annotated. The final publication graphic was sub-pixel antialiased (set antialias: 2) and depth-cued shadows were enabled.

### Theoretical target variant space calculations

To assess the accessibility of specific resistance alleles, we landscaped possible amino acid substitutions within the target gene *gyrA* QRDR. For this, we computationally generated all theoretical single-nucleotide variants (SNVs) for *A. baumannii* ATCC 17978 (GyrA amino acids 65–104^70^) and substituted each nucleotide position within the target window with all three alternative bases using the Biostrings package in R^71^. The resulting mutant DNA sequences were translated *in silico* to determine the corresponding amino acid residues. Variants were classified by their functional consequence as wild-type, synonymous, non-synonymous (missense), or nonsense (stop codon).

### Bacterial isolates and culturing

All bacterial strains in this study were derived from *Acinetobacter baumannii* ATCC 17978 (+AbaAL44). Unless stated otherwise, *A. baumannii* cells were plated on either Mueller Hinton II agar or LB agar (10g/L tryptone, 5g/L yeast extract, 5g/L NaCl, 15g/L agar) plates and liquid cultures were grown in Mueller Hinton II broth at 37°C, 200rpm.

### Fluctuation analysis and whole-genome sequencing of spontaneous mutants

Spontaneous mutation rates towards ciprofloxacin were determined using a Luria-Delbrück fluctuation assay in ATCC 17978^72^. For this, an overnight culture was diluted in fresh MHII to an OD600 = 5, i.e. approximately 1×10^10^ CFU/mL, serially diluted and day cultures of 2mL inoculated with ∼10^3^ CFUs. Day cultures were incubated for 5-6 hours (37°C, 220 rpm). To determine the total viable population size (N_t_), cultures were serially diluted in 1x PBS, and spot-plated on non-selective MHII agar. The remaining volume was plated onto large square MHII agar plates supplemented with ciprofloxacin at 2x, 4x, or 8x the wild-type MIC (1 ug/mL). CFUs were counted after overnight incubation at 35°C. Mutation rates were inferred using the Ma-Sandri-Sarkar maximum likelihood estimator method^73^. The final mutation rate (μ; i.e. number of mutations per cell per generation) was calculated as μ = m / N_t_. To identify target variants underlying the shifts in resistance, 239 spontaneous resistant colonies were isolated across concentrations and replicates of the fluctuation assay and prepared for whole-genome sequencing. Briefly, isolates were streaked twice on non-selective MHII agar and cells harvested in pellets. Genomic DNA extraction, library preparation and 2x 150bp sequencing on Illumina MiSeq were performed by Microsynth AG. Sequence variants compared to the ancestral, sensitive ATCC 17978 were called using a custom-build pipeline: Raw reads were trimmed using Trimmomatic^74^ v0.39 and aligned to the reference genome using Minimap2^75^ v2.26. Alignment BAM files were processed using Samtools^75^ v1.19, and genetic variants were called using FreeBayes^76^ v1.3.6, followed by quality filtering with BCFtools^77^ v1.19 and annotation using SnpEff^78^ v5.2 and Ensembl VEP^79^.

### Genome collection and target gene surveillance

*A. baumannii* genome assemblies were downloaded from NCBI (on 2025-10-15; n = 35,211) using the Entrez Direct API^80^. To establish the natural abundance of fluoroquinolone target gene missense mutations, target alleles were identified using a high-sensitivity TBLASTN search, and variant annotation was performed using SnpEff^78^ v5.2 against the ATCC 17978 reference genome (NCBI RefSeq assembly: GCF_902728005.1). The search was focused on target gene QRDRs (shifted from *E. coli* by -2 amino acids: *gyrA* (65–104^70^), *gyrB* (426–447^81^), *parC* (61–100^70^), and *parE* (458–458^82^)).

Additionally, a high-quality subset of genomes was then curated for later phylogenetic inference, based on (i) a coverage of ≥20x, (ii) ≤30 contigs, (iii) a BUSCO genome completeness score of ≥90% (inferred with BUSCO v5.8.2^83^ using the gammaproteobacteria_odb12 lineage dataset), (iv) an average nucleotide identity of ≥95% (confirmed using FastANI v1.34^84^ and *A. baumannii* ATCC 17978 as reference) and (v) no NCBI ExclFromRefSeq flags other than “from large multi-isolate project”, “derived from surveillance project”, “missing tRNA genes”, “missing rRNA genes”, “assembly spans repetitive elements”. To include assemblies with rare target variants but lower coverage or assembly completeness, we manually supplemented the collection with these assemblies leading to an overall collection size of 2,443 assemblies. All assemblies were uniformly annotated using Prokka v1.14.6^85^ (using the --genus Acinetobacter flag) for downstream analyses. Metadata were curated using NCBI BioSample host, host-disease, isolation-source, body-site, organism, and strain fields. Azure OpenAI GPT-4o (temperature 0) was used for preliminary closed-set metadata triage. Categories were assigned using a predefined deterministic hierarchical rule set, with evidence and unresolved records retained in an audit table (categories: human clinical, veterinary/animal, environmental, hospital-surface, laboratory/experimental, and unknown). For the origin-stratified QRDR analysis, *gyrA* missense substitutions at codons 65–104 were summarized within each origin category, and D70E variants were resolved by their underlying nucleotide change.

### Generation of isogenic *gyrA* mutant collection

GyrA QRDR target mutants were generated using two-step allelic exchange in ATCC 17978 (adapted from ^86^). Allelic exchange plasmids were based on the pVT77 suicide vector and contained the respective *gyrA* point mutations flanked by +/- ∼700 bp homology arms (synthesis by GenScript Biotech Corporation).

Allelic-exchange plasmids were transformed into the diaminopimelic acid (DAP)-auxotrophic *Escherichia coli* donor strain BW29427 using heat shock. Transformants were selected on selective LB agar (30 ug/mL kanamycin; 0.125 mg/mL DAP). For conjugation, overnight cultures of BW29427 (LB + kanamycin + DAP) and ATCC 17978 (MHII broth) were prepared. Donor and receiver (200 uL each) were combined, washed and resuspended in 50 uL LB. The cell mixture was spotted onto a nitrocellulose filter on selective MH agar (0.125 mg/mL DAP) and incubated at 30°C overnight. Subsequently, cells were harvested, resuspended in 1x PBS, and plated on selective LB agar (100 ug/mL sodium tellurite) to select for single-crossover integrants. After incubation (∼30 h at 37°C), 8–10 tellurite-resistant colonies per transformation were counter-selected in 0.5 mL LB broth with 1 mM isopropyl beta-D-1-thiogalactopyranoside (IPTG) for 1 h at 37°C and then streaked onto LB agar containing 200 ug/mL azidothymidine (AZT) and 1 mM IPTG. Thus, colonies of cells with excised plasmid could be picked the next days (∼30 per transformation) and were screened for successful mutagenesis by colony PCR and Sanger sequencing using primers gyrA_for_1 and gyrA_rev_2 (Supplementary Table 4; performed by Microsynth AG (Balgach, Switzerland). Confirmed mutants were purified on MH agar, validated for loss of vector backbone by selective plating and stocked in 20% glycerol at -80°C.

### Whole-genome validation of constructed D70E mutants

To validate genetic changes and genomic background of constructed ATCC 17978 *gyrA* D70E mutants (C>A and C>G) de novo Nanopore sequencing was performed by Microsynth AG (Balgach, Switzerland). Fastq reads from the sequenced isolates were aligned with minimap2 v2.24^75^ (-ax map-ont) to the *A. baumannii* ATCC 17978 assembly (GCF_902728005.1), used as the common coordinate and annotation reference, and BAM files were sorted and indexed with samtools v1.15^75^. The sequenced parental comparator (ROB01092) was aligned independently to the same reference, and loci uniquely callable in parent and mutants were included in parent-relative comparisons. Haploid variants were called with FreeBayes v1.3.6^76^ and retained at depth >20, QUAL >=30 and variant abundance level (VAL) >=0.80 before annotation with snpEff v5.1d^78^. Parent-different alleles were then compared across the three mutants; variants present in all three were classified as shared background, whereas only alleles restricted to one mutant or a two-mutant subset were considered potentially differentiating. Minority alleles were examined separately at 0.10 <= VAL <0.80 with strand, read-placement and independent pileup-count requirements. To assess coverage changes, *gyrA* depth was GC-corrected and normalized to callable single-copy windows within each sample, with mutant Nanopore and parental Illumina data normalized independently, and long-read breakpoint evidence was evaluated with Sniffles2 v2.6.3^87^.

### Antibiotic susceptibility testing (AST)

To quantify the level of resistance conferred by the *gyrA* QRDR mutations in ATCC 17978, minimum inhibitory concentrations (MICs) for ciprofloxacin were determined using the broth microdilution method according to CLSI guidelines^88^. In preparation, a 20x stock of ciprofloxacin (1,280 ug/mL) was prepared, serially diluted in distilled water and mixed with MHII broth to create the final 2x concentration series in round-bottom 96-well plates (50 uL/well; stored at -80°C until use). ATCC 17978 as well as quality control strains *Pseudomonas aeruginosa* ATCC 27853 and *Escherichia coli* ATCC 25922 were streaked on TSA and incubated at 37°C for 18-24h. Per colony (i.e. biological replicate), a bacterial suspension at 0.5 McFarland standard (1.5×10^8^ CFU/mL) was prepared by resuspending colonies in 2 mL sterile 0.85% NaCl and diluted in MHII (1:100) to reach a target density of 1×10^6^ CFU/mL. Compound plates were inoculated with bacterial suspensions (50uL/well) resulting in a final inoculum of 5×10^5^ CFU/mL. Each plate included sterility control (medium only) and no-drug growth control wells. After incubation at 35°C for 20 h, the MIC was determined by visual inspection as the lowest compound concentration that completely inhibited visible bacterial growth (i.e. MIC100).

### Bacterial growth kinetics and dose-response analysis

To measure how the *gyrA* QRDR mutations in ATCC 17978 impact growth and dose-response to ciprofloxacin, growth and dose-response experiments were conducted in tandem with the above AST. For both, the same bacterial inoculum prepared for AST was used to inoculate flat-bottom 96-well plates containing the two-fold ciprofloxacin dilution series in MHII medium in a randomized array. For 24 hours, plates were incubated at 37°C in a Biotek Synergy Neo2 multi-mode plate reader. Optical density (OD600) was measured every 20 minutes, directly after 30 s of orbital shaking. All assays were performed using four independent biological replicates, with three technical replicates for each. To determine drug-free fitness parameters (max. growth rate *r*, carrying capacity *k*, doubling time *t_gen*), growth curves of OD600 ∼ time from ciprofloxacin-free wells were analyzed using logistic growth modeling with the growthcurver package in R^89^. To quantify drug efficacy, dose-response curves were generated using the endpoint OD600 reading at 20 h (i.e. MIC scoring time point). Relative viability was calculated by normalizing OD600 under drug treatment to the corresponding drug-free control. Dose-response curves were generated using a four-parameter logistic model using the R package drc and the half-maximal inhibitory concentration (IC50) and Hill coefficients extracted accordingly.

### Time-kill kinetic assays

Time-kill kinetic assays comparing WT ATCC 17978 with *gyrA* QRDR mutants were performed using a method adapted from a persistence assay^31^. For pre-cultures, three colonies from MHII agar plates were inoculated into 3 mL of MHII broth in 14 mL round-bottom tubes and incubated overnight (37°C, 200 rpm). The following day, 8 uL overnight culture was sub-cultured in 8 mL fresh MHII broth in 50 mL tubes per biological replicate (37°C, 200 rpm). After 2.5 h (∼mid-logarithmic growth), an initial sample was collected (t=0h), and the remaining cultures were treated with a final concentration of 2 ug/mL ciprofloxacin (representing 2x the WT MIC). Further samples were collected at 0, 1, 3, 6, and 24 h post-treatment. Upon sample collection, bacterial suspensions were centrifuged at 10,000 x g for 2 min, washed once with sterile 1x PBS and resuspended in an equal volume of PBS. Resuspended samples were serially diluted 1:10 in PBS and serially plated on MHII agar for CFU quantification. After overnight incubation at 37°C, CFUs were counted and viable cell density calculated. This assay was performed with three independent biological replicates.

### Phylogenomics and ancestral state reconstruction

To derive the mutational history and drivers of *gyrA* evolution and to distinguish independent mutational emergence from clonal expansion, a high-resolution core-genome phylogeny was constructed. Because all-against-all orthology inference is computationally intractable for large genomic datasets, the 35,211 *A. baumannii* assemblies were first clustered to identify a non-redundant representative subset. This was achieved using the Microbial Agile Accurate SNP Typer (Maast v1.0.8 ^90^; min. SNP prevalence = 0.8) to define 118 “tag genomes” capturing the species’ core diversity. 11,411 orthologous groups for these tag genomes were subsequently inferred using the OMA Standalone tool v2.7.0^91,92^ using *A. baylyi* strain ADP1 (Accession: GCF_000046845.1) as an outgroup. Specific target variants from the overall genome collection were recursively mapped back into these groups using an fDOG-inspired^93,94^ targeted Hidden Markov Model (HMM)-based search. For each OG an HMM was built using hmmbuild from the HMMER suite^95^ v3.4 and used to identify the best hit against each of the proteomes from the genome collection (E-value cutoff = 1e-3, see also ^93^). Single-copy core orthogroups (n = 3,034) were aligned at the amino acid level using MAFFT^96^ v7.525 (--auto strategy), back-translated to codon alignments utilizing PAL2NAL^97^ v14.1, and trimmed using ClipKit^98^ v2.7.0 (settings: codon-aware and smartgap). A concatenated nucleotide supermatrix was used to infer a Maximum Likelihood starting tree using VeryFastTree^99^ v4.0.5 under a General Time Reversible (GTR) model. This was visualized in R using ggtree^100^. Ancestral State Reconstruction (ASR) was subsequently performed using IQ-TREE2^101^ with a GTR+G substitution model to map the evolutionary paths and quantify the discrete mutational events leading to extant *gyrA* alleles.

### Nucleotide substitution likelihood and neutral mutational propensity

To calculate the biochemical propensity and nucleotide substitution likelihood of specific evolutionary events, genome-wide 4-fold degenerate (wobble) sites were extracted from the codon alignments to establish a strictly neutral evolutionary supermatrix. A General Time Reversible (GTR) substitution matrix was inferred from this neutral subset using IQ-TREE2^101^ to infer nucleotide-specific substitution likelihoods and isolate intrinsic mutational biases (e.g., transition/transversion bias) from external selective pressures. Additionally, we calculated the transition-to-transversion (Ti/Tv) ratios to assess mutational bias.

### Multi-locus sequence type analysis

Genomes were assigned multi-locus sequence types (STs) as a high-level insight into clonal epidemiology^102–104^. Based on the Pasteur and Oxford seven-locus schemes, STs were inferred using mlst v2.11^105,106^ and the PubMLST.org website^107^. Genomes with >1 missing allele were discarded, ambiguous alleles were set to the first listed allele variant. Genomes with at least 6/7 loci detected were retained for subsequent analyses and consensus STs inferred for those with ambiguous assignments, based on the first-called alleles. To visualize genome similarities across the 7-locus allele profiles, we computed a pairwise Hamming distance matrix for all isolates and subsequently inferred an unrooted Neighbor-Joining tree, plotted using R packages ggtree^100^ and ggplot2^108^.

### Variant calling and evolutionary correlation across resistance targets

To determine if mutational substitution likelihood predicts resistance evolution across antibiotic classes, we profiled distinct antibiotic resistance-determining genes (*gyrA*, *pmrB*, *adeS*) across our curated *A. baumannii* genome collection. Variants were identified using targeted BLASTN against ATCC 17978 as above. Variants with at least three independent clinical observations were aggregated and mapped to the *A. baumannii* intrinsic substitution likelihood (P_sub_). Finally, to evaluate the predictive power of mutational bias, variants for each gene were ranked by global clinical abundance and the five most frequent were classified as dominant alleles. We ran regression analyses relating the P_sub_ and the log10-transformed abundance of the dominant alleles for each respective target.

### *gyrA* allele abundance versus substitution likelihood

Missense single-nucleotide variants in *gyrA* were extracted from the curated genome assembly collection and each was joined to its respective P_sub_. Abundance was taken as the number of genomes carrying that exact nucleotide change. The relationship was modeled by ordinary least squares as log10(allele count) ∼ P_sub, fitted independently within the QRDR, outside the QRDR, and across the full *gyrA* coding sequence (with and without the dominant S81L allele).

### Statistical analysis

To evaluate drug-free fitness costs, the relative carrying capacity (*k*) of each mutant was compared to the WT using a two-sided Student’s *t*-test. To correct for multiple hypothesis testing across the focal allele set, *P* values were adjusted using the Benjamini-Hochberg false discovery rate (FDR) method, with significance defined at an alpha level of 0.05. Finally, Negative Binomial regression generalized linear models (GLMs) were constructed to evaluate the relationship between observed allele abundance (across NCBI assemblies), the empirically determined IC50, drug-free growth rate (*r*), and the nucleotide substitution likelihood calculated above (MASS package^109^ in R, utilizing a log-link function to handle overdispersed genomic count data). Allelic abundance was modeled as a linear combination of standardized (Z-scored) biological predictors: relative resistance (IC50), relative fitness (growth rate *r*), and nucleotide substitution likelihood. Variance Inflation Factors (VIF) were calculated prior to modeling to verify the absence of multicollinearity among these predictors, ensuring the independent contribution of each variable could be accurately assessed.

### Pharmacokinetic and Dynamic Killing Framework

To translate *in vitro* mutant phenotypes into clinical outcomes, we modeled systemic ciprofloxacin concentrations (*C(t)*) using a one-compartment oral pharmacokinetic framework with first-order absorption (*k_a_*) and first-order linear elimination (*k_el_*), parameterized by apparent volume of distribution (*V*/*F*) and systemic clearance (*CL*/*F*), and twice daily dosing with 500mg ciprofloxacin, as FDA-recommended^34^. We considered dosing with ciprofloxacin hydrochloride (MW = 385.8 g/mol). For each virtual patient, the unbound peak concentration (*C_max_*), time to peak (*t_max_*) and elimination rate constant (*k_el_*) were drawn from truncated log-normal distributions (Supplementary Table 5; ); *k_a_* was then identified per patient by grid search over 1.49–2.80 h⁻¹ (step 0.05 h⁻¹) as the value whose simulated t_max best matched the sampled t_max, and the simulated drug amount was scaled to reproduce the sampled *C_max_*. The *V/F* and *CL/F* were therefore not sampled directly; they are implied by the per-patient scaling factor (≡ 1/(*V/F*)) and *k_el_* (*CL/F* = *k_el_/F*). Virtual cohorts (*n*=100,000 per population) were generated via vectorized rejection sampling from log-normal distributions to account for physiological variability across healthy, renally-impaired, and low-penetration populations (Supplementary Table 5; ^33^). For each virtual patient, the instantaneous drug effect (*E(t)*) on specific *gyrA* alleles was derived using the Hill equation: 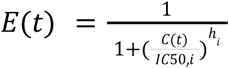 where IC50 and h represent allele-specific pharmacodynamic parameters. For synonymous D70E variants, codon-specific parameters were calculated based on the *in vitro* data.To bridge continuous pharmacokinetic profiles with the stochastic evolutionary model, we implemented a dynamic kill rate (α*_i_* (*t*)) and a comparative time-averaged effective kill rate (α*_i, ave_*) calculated over a 14-day window: 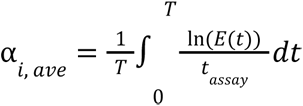. The choice of *t _assay_* = 1ℎ was empirically anchored to our experimental time-kill kinetics (Supplementary Data 3b), in which ciprofloxacin exhibited rapid, concentration-dependent bactericidal action where the primary multi-log reduction in viable cells occurred within the first hour of drug exposure.

### Stochastic Evolutionary Dynamics

Bacterial population dynamics of wild-type (WT) and mutant alleles were simulated using an adaptive tau-leaping algorithm to capture probabilistic rare-event mutations and extinctions based on ^5^. The model integrates host ecology by decoupling net growth into absolute birth (b_i_) and death (d_i_) rates under a fixed *in vivo* spatial turnover constraint (T=0.9). In the absence of significant growth costs of the *gyrA* QRDR mutations in the reconstructed strains (Fig. 1d,e), we modeled uniform bacterial growth rates based on the measured wild-type maximum growth rate *in vitro*. To align *in vitro* growth kinetics with clinical disease progression, we applied an *in vivo* translation scaler (γ = 0.25) to all growth rates, effectively calibrating the model to replicate observed doubling times in human pneumonia^67,110^ as well as a carrying capacity of 10^10^ cells^111,112^. Host-mediated innate immune clearance impairment in neutropenic patients was parameterized as δ_human_ = 0.038 h^−1^. This parameter was derived from early-passage *A. baumannii* ATCC 17978 lung clearance kinetics in mice prior to resistance selection (Passage 2 in ^31^), where lung burdens at 27h post-infection in cyclophosphamide-treated neutropenic mice diverged by 1.789 log_10_ CFU from immunocompetent controls (geometric mean CFU = 1.35 × 10^5^ CFU; log_10_ = 5.131), defining a murine clearance differential of: 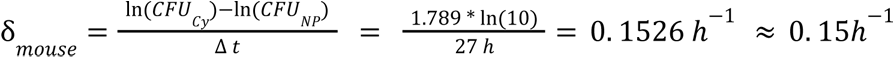. To translate this preclinical rate to human pulmonary physiology, δ_k_mouse_ was scaled using an *in vivo* translation factor of 0.25, derived from quarter-power allometric scaling of physiological rate constants (*k* ∝ *M*^−0.25^) and pulmonary clearance turnover between standard reference rodent and human scales^113,114^, yielding: δ*_mouse_* = δ*_mouse_* * 0. 25 = 0. 03815 ≈ 0. 038 ℎ . In simulated neutropenic cohorts, δ*_ℎuman_* was subtracted from the baseline bacterial death rate to model the selective loss of neutrophil-mediated bactericidal activity. While growing, simulated bacteria could become resistant, i.e. mutate into any of the included *gyrA* QRDR alleles, given the empirically-determined resistance rate, *μ*, of 1×10^−7^ (Supplementary Table 6;^20,115,116^). Mutations arise during replication as a birth-coupled event yielding one mutant daughter, with the wild-type parent retained. Simulations were performed in R v4.2.0 using the deSolve^117^ and parallel packages.

### Model sensitivity and fit evaluation

Parameter sensitivity and model fit robustness were evaluated by scoring simulated post-treatment mutant reservoir distributions against empirical isolate counts of *A. baumannii gyrA* quinolone resistance-determining region (QRDR) variants in the NCBI-based genome collection. We implemented a multinomial log-likelihood objective score, 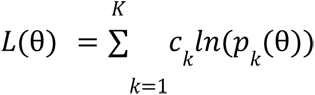, where *c_k_* represents the observed genomic count of allele *k*, and *p_k_* (θ) represents the relative proportion of allele *k* across rebounding virtual patients. A one-at-a-time sensitivity grid was evaluated around the empirical baseline parameter set (resistance rate mu = 10^−7^, a turnover rate of T = 0.9, and unit scalers for drug kill rate, relative growth rate, and initial inoculum size; see Supplementary Table 6) across 100,000 simulated virtual patients per parameter state. Single-parameter perturbations were evaluated to quantify relative changes in log-likelihood fit (Δ*L*) and evaluated whether clinical rebounds were observed. A trajectory was classified as a rebound only if it satisfied three conditions concurrently: (1) a ≥ 2-log10 bacterial expansion following the post-treatment nadir^20,43^ (with a special case allowing a peak ≥10^2^ cells if complete transient sterilization, nadir = 0 cells was achieved), (2) attainment of a peak absolute mutant burden ≥10^4^ cells, and (3) a final total surviving bacterial population ≥10^3^ cells at the end of the simulation window. Profiles failing any of these criteria were classified as successfully cleared. We confirmed clinical realism by checking that the optimized parameter set led to a simulated rebound rate between 1% and 5%^44,45^ in a cohort of healthy patients. The evolutionary success of resistance alleles was thus quantified as the resistance reservoir potential (Area Under the Curve (AUC,^44^), of a given mutant abundance post-treatment.

### Comparative Assessment of Clinical Cohorts and Evolutionary Drivers

To isolate the determinants of resistance emergence, we performed a series of comparative simulations and parameter ablation studies. To evaluate the relative weight of biological drivers, we compared the best-fitting mechanistic model against “null” scenarios where specific parameters, namely the substitution likelihood (*P_sub,i_*), or drug-induced kill rate (α ) were set to be uniform across all alleles, in the case of α*_i_*, to the S81L-specific α. In the clinical cohort analysis, we generated 100,000 baseline healthy profiles and systematically applied physiological modifiers (see Supplementary Table 5) to simulate renally impaired, and low-penetration scenarios (0.2), ensuring that observed differences in mutant success were attributable strictly to PK/PD variance. Population-scale results were aggregated by calculating the mean relative abundance and standard error of each *gyrA* allele across rebounding patients. The superiority of the dynamic model over the static time-averaged approach was quantified using a multinomial log-likelihood comparison of the simulated mutant reservoir composition against empirical clinical isolate frequencies.

## Acknowledgements

We thank M. Leipner, S. Gartenmann, F. Dey, J. Payne, M. Lobritz for constructive discussions and support and A. Praetor and N. Alonso for external alliance management. This project was funded by a Roche Postdoctoral Fellowship awarded to N. O.

## Author contributions

N.O., A.T., A.J.M.-B.., and J.P. conceptualized the study; N.O., A.R. and Q.N. performed experiments; N.O and D.D. developed phylogenetic analysis pipeline; N.O. executed bioinformatic analyses; N.O. and J. P. developed simulation model; K.B. and A.J.M.-B. provided resources and sequencing support; A.T., J.P., K.B. and A.J.M.-B. provided supervision; N.O. analyzed the data and wrote the original draft of the manuscript; all authors reviewed and approved the final version for submission.

## Ethics declaration

not applicable.

## Data Availability Statement

The raw data and code files used for this study are deposited in a repository (DOI: 10.5281/zenodo.24308153) repository and will be made available to editors and reviewers.

## Code Availability Statement

The code files used for this study are deposited in a Zenodo repository (DOI: 10.5281/zenodo.24308157) repository and will be made available to editors and reviewers.

---

**Supplementary Data 1.**
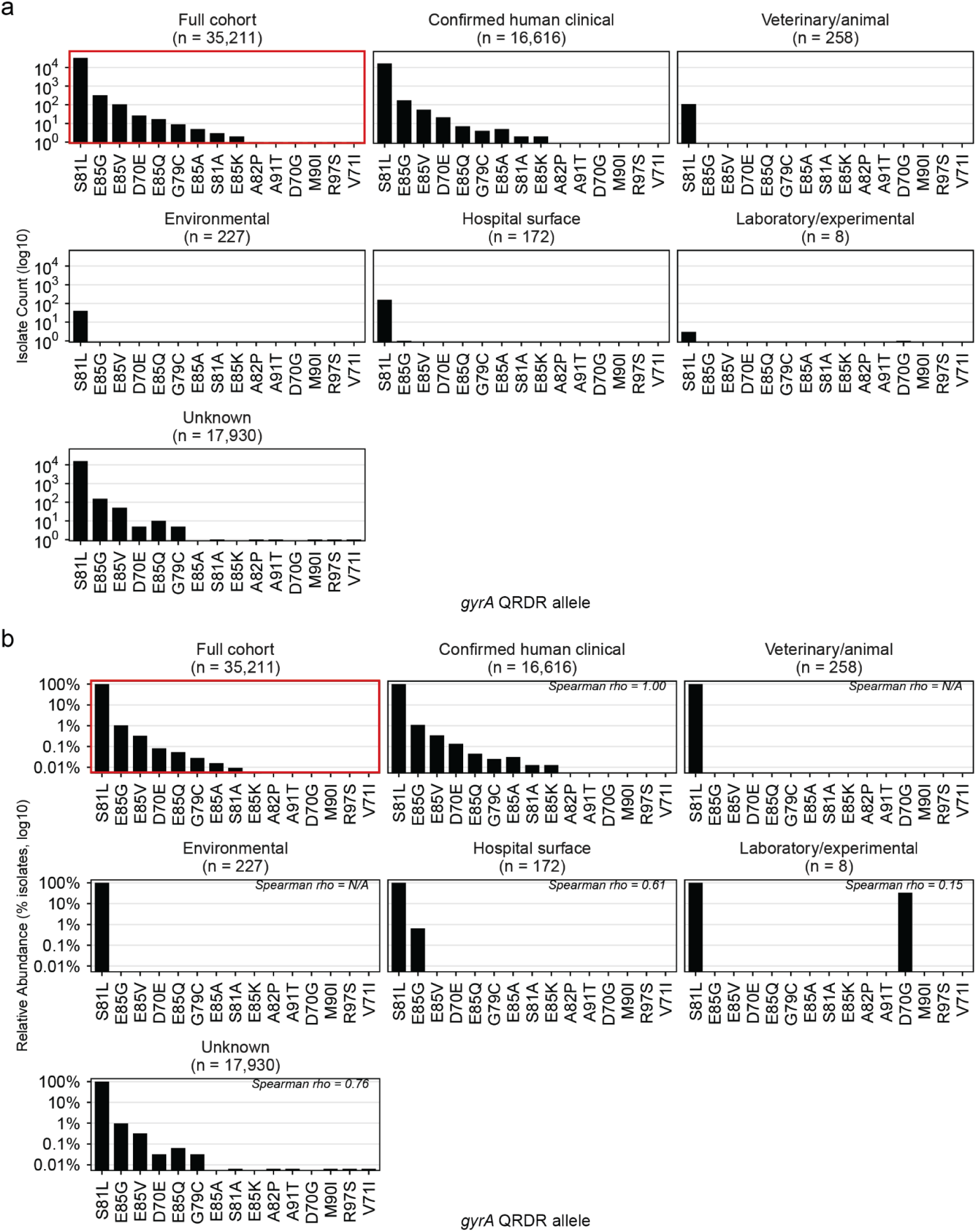
Origin-stratified distribution of *gyrA* quinolone-resistance determining region (QRDR) substitutions across the NCBI *A. baumannii* genome collection. **a**,Absolute abundance of detected *gyrA* QRDR missense substitution in the full cohort and in confirmed human clinical, veterinary/animal, environmental, hospital-surface, laboratory/experimental, and unknown source subsets shown on a log10 scale. **b**, Relative composition of observed substitutions among QRDR-substitution-bearing isolates in each subset. The most common variant, S81L, is shown in red, all other variants in black. For subsets with more than a single *gyrA* QRDR mutation, Spearman rank correlations were used to compare the per-allele relative frequency profile of a subset with that of the full collection.

**Supplementary Data 2.**
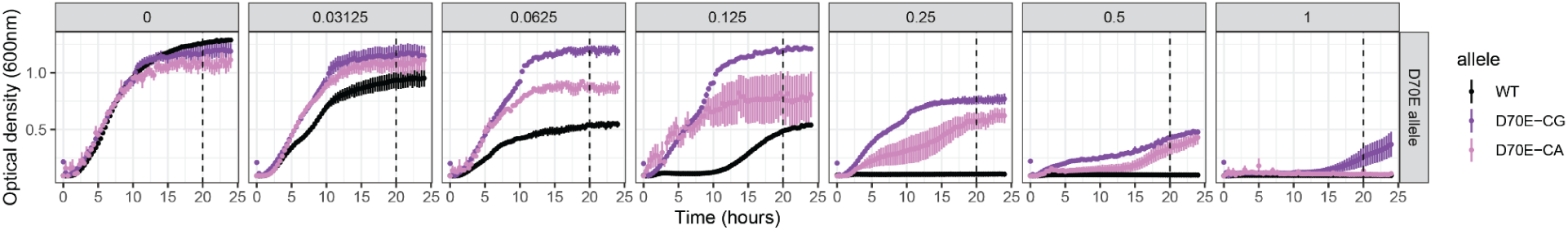
Growth kinetics of *A. baumannii* ATCC 17978 WT and synonymous *gyrA* D70E mutants across different ciprofloxacin concentrations. Time-resolved bacterial growth dynamics (OD600 over 24 h) measured across a two-fold ciprofloxacin concentration gradient for susceptible wild-type (WT) *A. baumannii* ATCC 17978 alongside two isogenic *gyrA* QRDR variants encoding the identical D70E amino acid substitution:C>A (D70E-CA; light purple) and C>G (D70E-CG; dark purple). Average OD600 (-/-s.e.m) shown for 4 independent biological replicates (each with 3 technical replicates).

**Supplementary Data 3.**
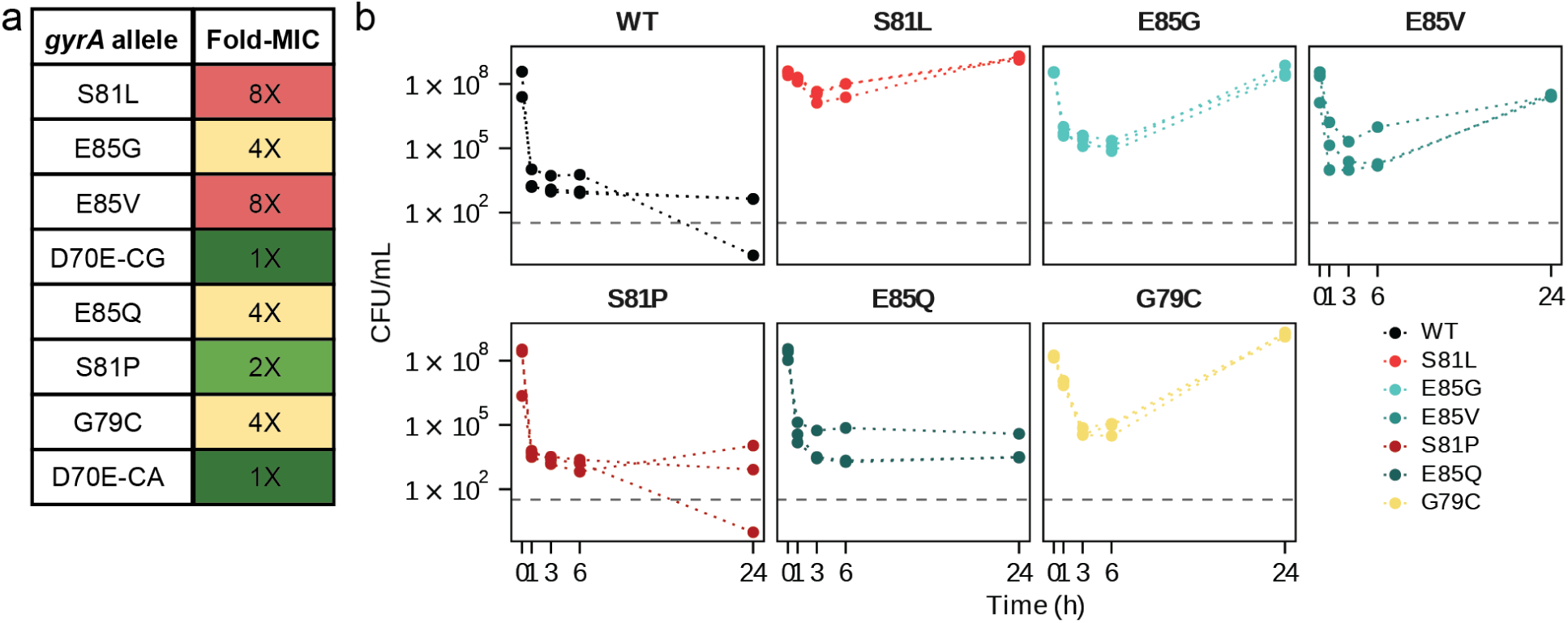
Extended resistance characterization of isogenic *gyrA* QRDR mutants in ATCC 17978. **a**,Fold-change in minimum inhibitory concentrations (MICs) compared to susceptible WT ATCC 17978, with an MIC of 1ug/mL ciprofloxacin in MHII broth. Mode MICs of 5 biological replicates are shown. **b,** Time-kill analysis of bacterial isolates exposed to 2 *μ*g/mL ciprofloxacin in MHII broth. Curves show exponential fit (with 95% CI) of 3 biol. replicates (data points). Across all strains, the steepest bactericidal clearance occurs during the initial 0–1 h interval (Δlog_10_*CFU*/ℎ), which served as the empirical basis for anchoring the dynamic killing rate normalization (*t_assay_* = 1ℎ) in the evolutionary model.

**Supplementary Data 4.**
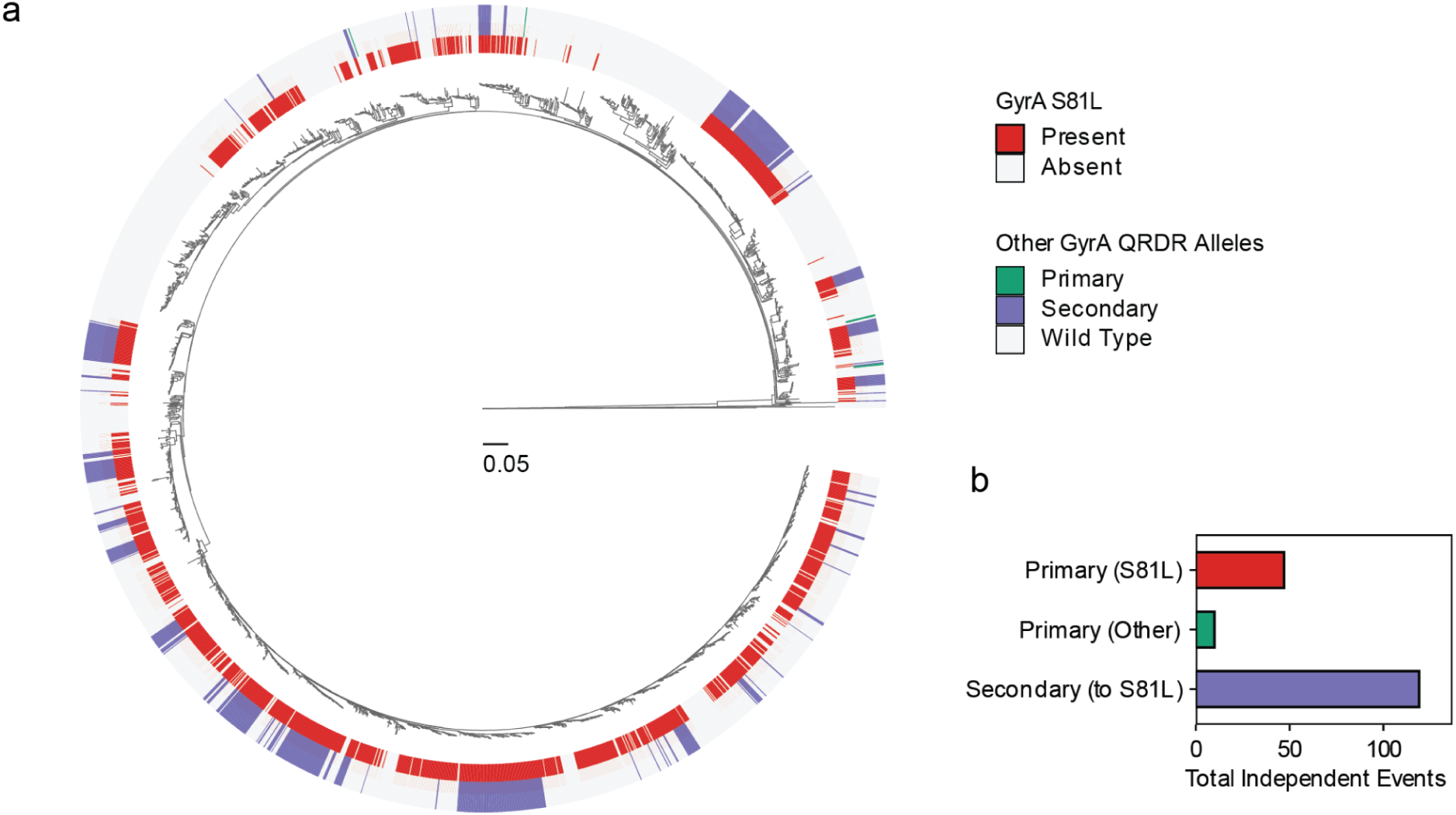
Extended phylogenetic analysis of *gyrA* QRDR evolution. **a**,Core genome maximum-likelihood phylogeny of 2,443 high-quality *A. baumannii* genomes, rooted to the outgroup *A. baylyi* ADP1. The inner ring designates the presence (red) or absence (light grey) of the primary driver mutation S81L. The outer ring displays the evolutionary context of other *gyrA* QRDR alleles, classified as primary mutations (emerging independently of S81L; teal) or secondary mutations (co-occurring with S81L; light blue). The scale bar represents nucleotide substitutions per site. **b**, Absolute abundance of independent mutational events inferred via ancestral state reconstruction, quantifying the emergence of the S81L driver versus other primary and secondary alleles.

**Supplementary Data 5.**
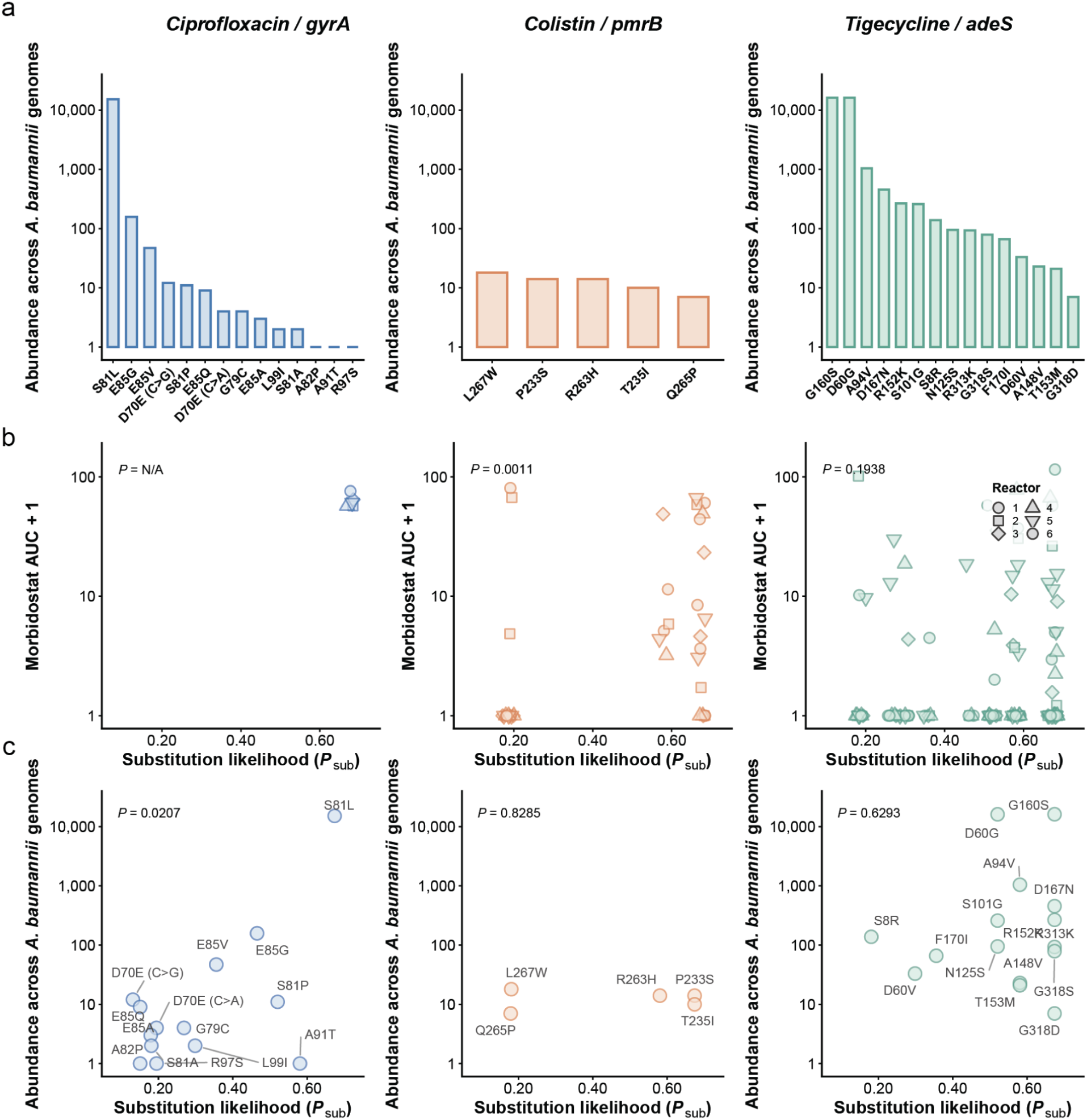
Multi-scale mapping of evolutionary landscapes reveals a progressive decay of mutational signaling across a continuum of target constraints. **a**,Epidemiological baseline of unique resistance alleles in terms of unique variant counts observed across public *A. baumannii* genome assemblies. **b,** Laboratory evolution dynamics captured across independent parallel morbidostat bioreactor runs (N = 6 independent replicates per antibiotic, indexed by geometric shapes, from Zlamal 2021, Kent 2025). Observed variant selection magnitudes (Area Under the Curve; Morbidostat (AUC+1)) are plotted against nucleotide substitution likelihoods. Solid markers track individual uncollapsed replicates, i.e. bioreactors. Statistical metrics denote two-sided Spearman’s rank correlation coefficients (⍴) and corresponding P-values; no correlation was performed for *gyrA*, due to the emergence of only a single variant (S81L). **c,** Correlation between intrinsic nucleotide substitution likelihood and long-term abundance across genome assemblies. Unique amino acid substitutions are labeled. Statistical values indicate two-sided Spearman’s rank correlation coefficients and corresponding P-values.

**Supplementary Data 6.**
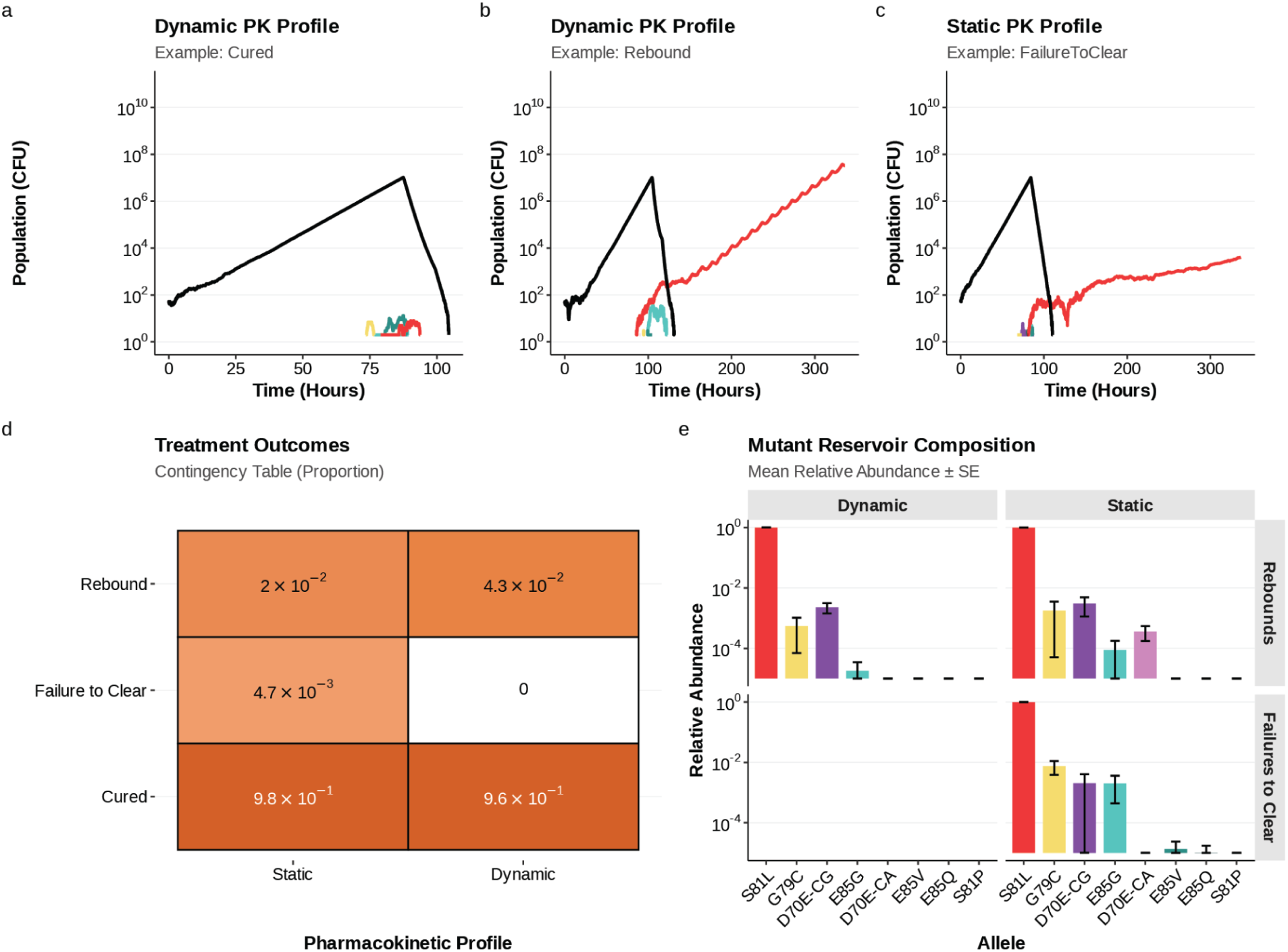
Comparison of dynamic vs. static drug kill rate (α) implementations in the stochastic evolutionary model. **a-c**, Representative *in silico* bacterial population trajectories (CFU) over time comparing dynamic vs. static pharmacokinetic profiles: **a**, Dynamic PK profile resulting in complete clinical cure, showing rapid wild-type clearance (WT, black line) and transient, non-establishing mutant subpopulations (colored lines). b, Dynamic PK profile resulting in clinical relapse, where instantaneous drug concentration fluctuations below mutant MICs during PK troughs promote characteristic “sawtooth” population expansion dominated by the primary driver allele S81L (red line). **c**, Static PK profile (time-averaged kill rate α) resulting in persistent failure to clear, where constant drug pressure leads to continuous, monotonic mutant expansion lacking PK trough dynamics. **d**, Contingency heatmap comparing population-scale treatment outcome proportions (rebound, failure to clear, cured) between static and dynamic model architectures across N = 100,000 virtual patients. **e**, Post-treatment mutant resistance reservoir composition (mean relative abundance s.e.) stratified by clinical outcome (rebounds vs. failures to clear) and PK profile.

**Supplementary Data 7.**
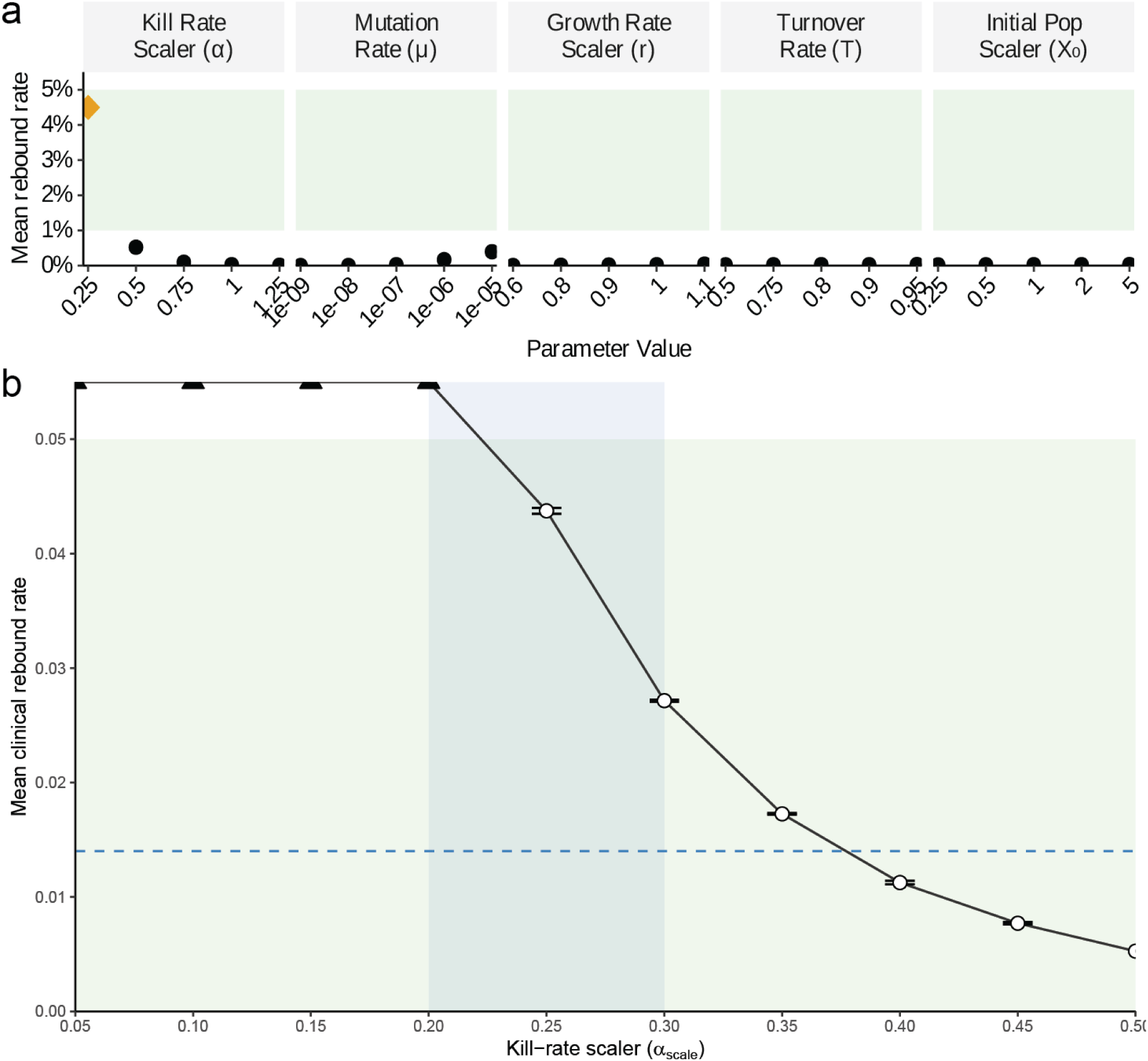
Sensitivity analysis and fine local sweep of the *in vivo* kill-rate scaler (α). **a**,One-at-a-time (OAT) sensitivity analysis evaluating the impact of individual parameter variations - kill-rate scaler (**α**), global mutation rate (*μ*), growth-rate scaler (*r*), turnover rate (T), and initial population scaler (X0) - around the baseline model configuration. The horizontal green shaded band denotes the acceptable clinical rebound window (0-5%; horizontal green shaded band). Points represent mean rebound rates across simulation replicates (N = 5 independent replicates per parameter value, up to 100,000 virtual patients per replicate); error bars denote 1 s.e.m. . The orange diamond highlights the selected *in vivo* kill-rate scaler value (**α** = 0.25). b, Fine local sensitivity sweep of the kill-rate scaler (**α**) evaluated from 0.05 to 0.50 in increments of 0.05. White circles represent parameter values yielding rebound rates within the target window, with error bars denoting 1 s.e.m. across 5 independent replicates (up to 100,000 virtual patients per replicate, unless rebound rate is > 5%, which abrograted simulations to save computation); black triangles represent parameter configurations exceeding the 5% rebound threshold, capped at the 0.055 upper plot boundary due to early stopping.

**Supplementary Data 8.**
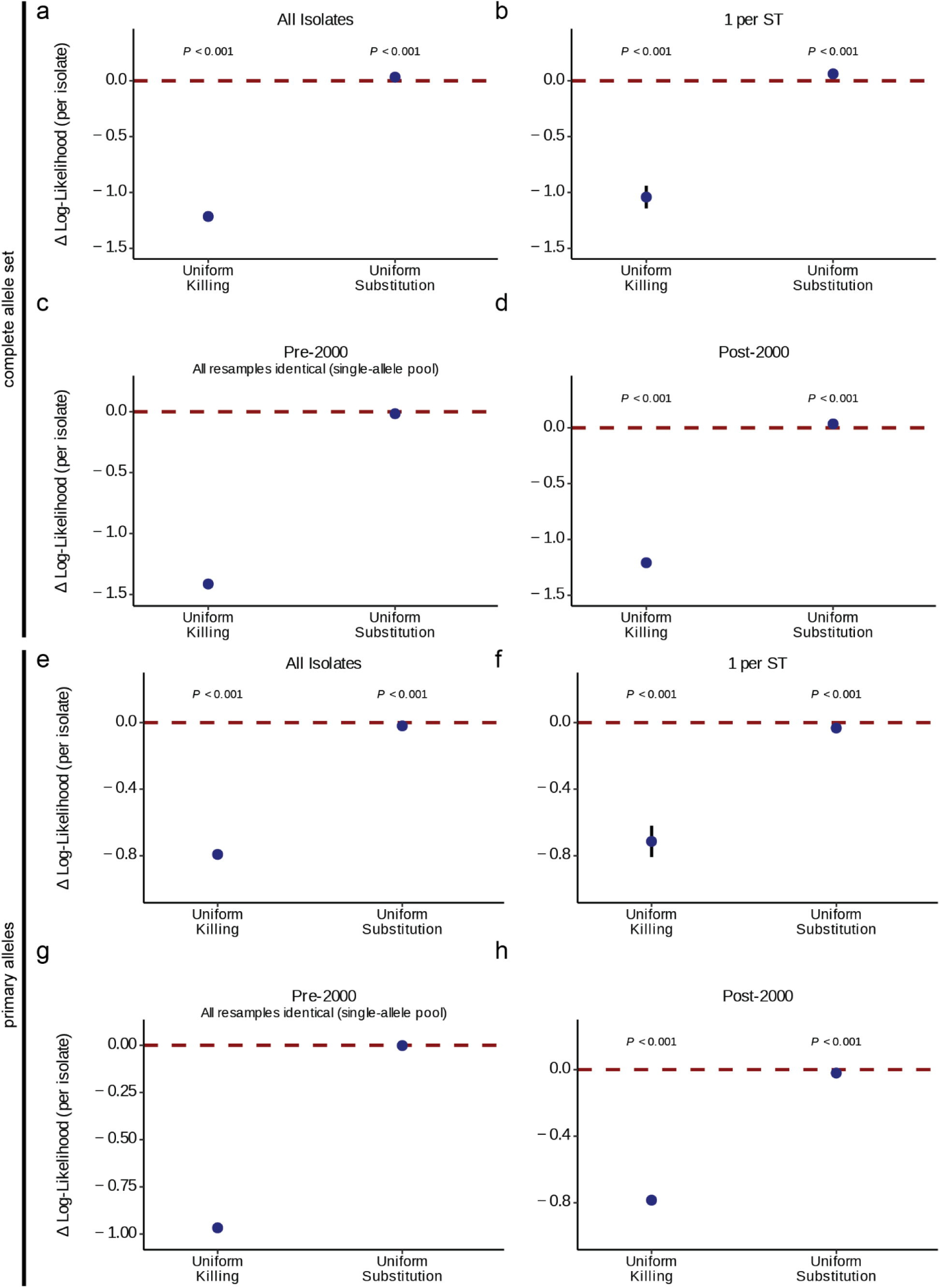
Bootstrapping analysis verifying model fit robustness against phylogenetic and temporal collection biases. **a-h**, Assessment of model fit performance (change in log-likelihood per isolate relative to the full baseline model) across 1,000 bootstrap iterations evaluating distinct genomic subsampling scenarios: **a**, Random bootstrap resampling across *Acinetobacter baumannii* NCBI genome assemblies (N = 35,211). **b**, Phylogenetically de-replicated subsetting by focusing on single randomly selected isolates per Pasteur Multi-Locus Sequence Type to remove ST2 clonal bias, clonal expansion and outbreak surveillance biases. Note that this removes rare alleles observations from some ST subsets. **c, d,** Temporal partitioning comparing isolates collected in early vs. late epidemiological eras: c, pre-2000 vs. d, post-2000, aligning with early sequencing surrounding the market introduction of ciprofloxacin for the pre-2000 cohort. Note that the pre-2000 cohort includes 139 genomes, with 77 mutants, all of which are S81L, thus leading to s.d. = 0 by definition. Across panels, ablated model scenarios are displayed along the horizontal *x*-axis and per-isolate Δ log-likelihood changes along the vertical *y*-axis. Point show mean paired Δ log-likelihood values across 1,000 independent bootstrap iterations and vertical error bars (+/- 1 s.d.). Horizontal red dashed lines mark the benchmark performance of the baseline model (Δ = 0). FDR-corrected *P*-values from paired two-sided Wilcoxon signed-rank tests across bootstrap replicates compare each ablated model scenario (substitution likelihood, drug-induced killing) against the baseline model. **e-h**, Show ablation analysis results based on simulations with primary *gyrA* QRDR alleles (S81L, E85G, E85V, G79C) only.

**Supplementary Data 9.**
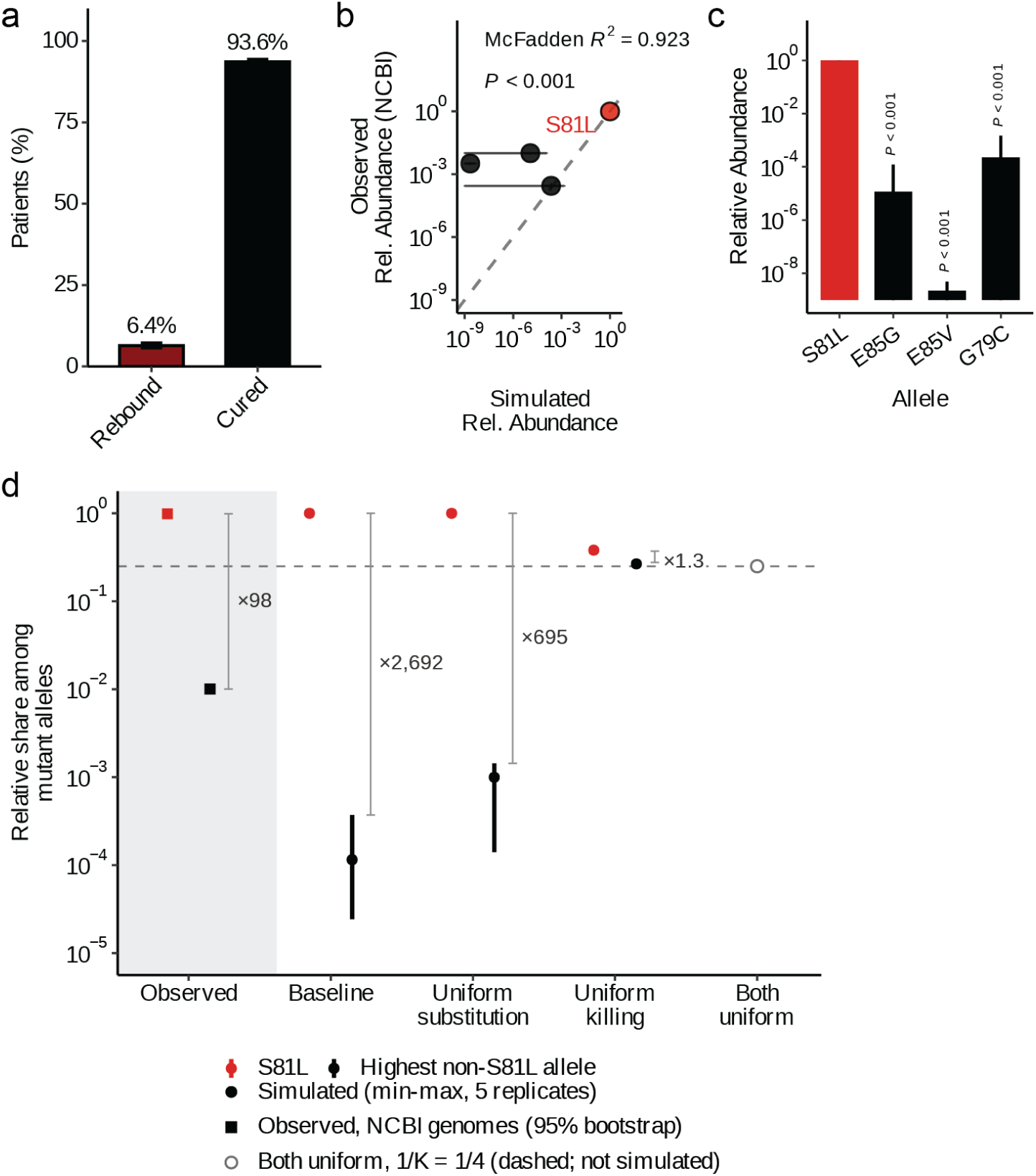
Mechanistic multi-scale modeling focusing on primary *gyrA* QRDR allele prevalence. **a**,Simulated clinical outcome frequencies for the healthy baseline cohort (*N* = 100,000 virtual patients). Bar heights indicate the median percentage of patients classified as rebound or cured across simulation batches; error bars denote ±1 s.d. **b**, Concordance between simulated relative allele abundance in the post-treatment resistant reservoir and observed global prevalence from NCBI genomes. Colored points represent simulated *gyrA* mutant alleles; the dashed line indicates 1:1 identity. Goodness-of-fit is evaluated via multinomial log-likelihood estimation against a uniform null model (McFadden’s R^2^ and Likelihood Ratio Test P-value); error bars show inter-patient population dispersion (+/- 1 s.d.) across simulation batches. **c,** Predicted relative abundance of each primary *gyrA* QRDR allele in the resistant reservoir under the baseline model (log10-scale). Error bars show inter-patient population dispersion (+/- 1 s.d.) across simulation batches. The statistical difference between S81L abundance vs. the each other allele is indicated by FDR-corrected *P*-values from two-sided Mann–Whitney U tests. **d,** Model component attribution by parameter ablation. For the baseline model and for each ablated scenario, points show the share of the simulated post-treatment resistant reservoir held by S81L (red) and by the most abundant non-S81L allele (black, runner-up depending on the simulation), alongside the corresponding observed shares among NCBI genomes (filled squares, shaded panel at left). Allele-specific drug-induced kill rates (“Uniform kill.”, set to α_S81L_) or allele-specific nucleotide substitution likelihoods (“Uniform sub.”) are replaced by a single value shared across alleles. Error bars denote +/-1 s.d. across five independent replicate simulations of 100,000 virtual patients each. Grey brackets and annotations give the dominance margin between S81L and the runner-up allele, defined conservatively as the minimum S81L share across replicates divided by the maximum share attained by any non-S81L allele in any replicate. The dashed horizontal line and open circle mark the chance expectation 1/K = 0.125 for K = 4 exchangeable alleles, i.e. uniform allele proportions.

## Supplementary Tables

**Supplementary Table 1.** Literature overview of *gyrA* QRDR variants.

**Supplementary Table 2.** Genomic validation of constructed D70E nucleotide variants.

**Supplementary Table 3.** Relationship of nucleotide substitution likelihood on *gyrA* SNP mutations.

**Supplementary Table 4.** Primers and plasmids used in this study.

**Supplementary Table 5.** Parameters and variables used in pharmacokinetic modeling.

**Supplementary Table 6.** Parameters and variables used in stochastic evolutionary simulations.

